# SOP-MULTI: A self-organized polymer based coarse-grained model for multi-domain and intrinsically disordered proteins with conformation ensemble consistent with experimental scattering data

**DOI:** 10.1101/2024.04.29.591764

**Authors:** Krishnakanth Baratam, Anand Srivastava

## Abstract

Multidomain proteins with long flexible linkers and full-length intrinsically disordered proteins (IDPs) are best defined as an ensemble of conformations rather than a single structure. Determining high-resolution ensemble structures of such proteins poses various challenges using tools from experimental structural biophysics. Integrative approaches combining available low-resolution ensemble-averaged experimental data and *in silico* biomolecular reconstructions are now often used for the purpose. However, an exhaustive Boltzmann weighted conformation sampling for large proteins, especially for ones where both the folded and disordered domains exist in the same polypeptide chain, remains a challenge. In this work, we present a 2-site per amino-acid resolution SOP-MULTI force field for simulating coarse-grained models of multidomain proteins. SOP-MULTI combines two well-established self-organized polymer (SOP) models —: (i) SOP-SC models for folded systems and (ii) SOP-IDP for IDPs. For the SOP-MULTI, we train the cross-interaction terms between the beads belonging to the folded and disordered regions to generate experimentally-consistent conformation ensembles for full-length multi-domain proteins such as hnRNPA1, TDP-43, G3BP1, hGHR-ECD, TIA1, HIV-1 Gag, Poly-Ubiquitin and FUS. When back-mapped to all-atom resolution, SOP-MULTI trajectories faithfully recapitulate the scattering data over the range of the reciprocal space. We also show that individual folded domains preserve native contacts with respect to solved folded structures, and root mean square fluctuations of residues in folded domains match those obtained from all-atom molecular dynamics simulations trajectories of the same folded systems. SOP-MULTI Force Field is made available as a LAMMPS-compatible user package along with setup codes for generating the required files for any full-length protein with folded and disordered regions.

## 1 Introduction

The advent of AlphaFold has unleashed a sudden increase in the availability of protein structural models.^1,2^ However, the availability of a static structural model of the protein alone is insufficient to understand the functional implications of the protein. The biological aspects of the given protein are appreciated better by the ensemble it constitutes in the cellular environment.^3^ The heterogeneity in the ensemble is determined by the nature of the protein and its environment. A well-folded protein exhibits small conformational fluctuations around its mean native structure. On the other hand, an intrinsically disordered protein (IDP) lives in a dynamic equilibrium among its various possible conformations. Similar heterogeneity is also observed in multidomain proteins (MDPs) comprising multiple folded domains connected by flexible linkers or terminal disordered regions. ^4–6^ Despite their vast conformational heterogeneity, the MDPs and IDPs are physiologically very important and the wide prevalence of IDPs and MDPs across proteomes of various species indicate their functional role in various spatially and temporally coordinated events such as catalysis, signalling, gene expression regulation and other cellular processes. ^7–10^

IDPs and MDPs are also implicated in several pathological conditions of human diseased states.^11,12^ Transactive response DNA-binding protein (TDP-43), hnRNPA1, and human fused in sarcoma/translocated in liposarcoma (FUS/TLS) are some well-known examples of RNA- and DNA-binding proteins with multiple folded domains and IDRs.^13^ Mutations discovered in the sequences of TDP-43 and FUS are known to increase the propensity of formation of pathological aggregates of these proteins^14^ leading to disease states such as Amyotrophic lateral sclerosis (ALS) and frontotemporal lobar degeneration (FTLD). Similarily, hnRNPA1 has many roles, including gene regulation, translation, splicing, export via nuclear pores and transcription of cellular and viral transcripts. In addition to mRNA biogenesis, hnRNPA1 also performs a variety of other tasks, such as microRNA processing, telomere preservation, and transcription factor activity modulation. ^15^ Hybrid proteins such as these, with both folded and disordered regions, function by a tuned molecular interplay between their folded domains and IDR. The physiological relevance of these proteins makes them an attractive target for discovering new classes of drugs that can modulate the formation of pathological aggregates. ^16^ One of the major area of research that has gained traction is related to understanding how the molecular grammar is encoded in the sequence of these proteins^17^ as well as to decipher the interactions at play by integrating molecular simulations and experimental observations. ^18^

Larger IDPs^19^ and MDPs^20^ pose challenges in exhaustive sampling of their conformational space due to their flexibility and their size. Molecular simulations with atomistic resolution are either not feasible or pose convergence issues with current computational resources. Coarse-grained models are a practical way out in reaching such lengths and time scales. A wide variety of coarse-grained models of IDPs have recently been developed in this regard. Kim-Hummer,^21^ HPS-KR,^22^ Bead-Necklace model,^23^ FB-HPS,^24^ HPS-URRY,^25^ SOP-IDP,^26–28^ MPIPI,^29^ CALVADOS,^30^ Martini3^31^ are some recent models that have been used with varying degree of success in recent past to model IDP-related systems and get deeper molecular-level insights behind the underlying biophysical processes.^32,33^ Unlike pure IDP CGFFs, the multidomain protein CGFF are very limited in number and need different parameterization prescription since they carry both the folded and disordered regions. In general, folded regions are maintained by either considering it as a rigid body using a scaled set of force field parameters^21,34,35^ or by applying restraints.^36,37^ In the field, there is an immediate need to extend the IDP models for multidomain proteins, which share challenges similar to those of IDPs. Of the several IDP CGFF models, CALVADOS^38^ and Martini^39^ have now extended their models to accommodate both IDPs and MDPs. This article focuses on extending the self-organized polymer (SOP) class of CGFF protein models to MDPs. The Self Organized Polymer model has two separate formulations for folded proteins^40^(SOP-SC) and pure IDPs^26–28^(SOP-IDP), thus making it an attractive model for the development of a hybrid model for MDPs and folded proteins with long IDRs. Also, with a 2-site per amino acid representation, the model works very well with all-atom reconstruction algorithms and is very useful in extraction of high-resolution conformation ensemble of these flexible proteins from their coarse-grained representations.

The SOP-SC model has been used extensively to study allosteric transitions, ^41^ folding and unfolding pathways,^42,43^ denaturant dependent folding, ^44–46^ energy landscapes^47^ and folding kinetics^48^ of well-folded proteins. On the other hand, the SOP-IDP model was developed relatively recently and has shown promising results in capturing ensembles of IDPs that are consistent with available experimental data^26–28,49,50^ and providing insights into IDPs involved in condensate formation^51^ and those with propensity to aggregate. ^52,53^ In this work, we harness these two models to develop a hybrid model for handling multidomain proteins. We make this model available in the form of SOP-MULTI, a package of C++ codes that can be patched to LAMMPS^54^ simulation engine. We also provide the supporting Python codes that facilitate the setup of input files for a given IDP or multidomain protein. ^55–57^ We show the application of our code on ten full-length multidomain proteins. For backward compatibility, we also tested the code on twenty-two pure IDPs that were reported previously.^28^

The rest of the paper is organized as follows. The details of the hybrid model and its force field, along with a description of the analyses used, are explained in the Material and Methods section below. In the Results and Discussion section following the Material and Methods section, we report the observations from this model in light of previously reported experimental and simulation data on several MDP and IDP systems. In the last section of the manuscript, we provide a conclusion summary and also discuss the scope and limitations of the model. The paper comes with two companion documents, the first one being the supporting information files with corroborating data for the manuscript (SI-1) and the second document is the the user-manual for SOP-MULTI for both the users of the package as well for future developers (SI-2).

## 2 Material and Methods

### 2.1 SOP-MULTI model

SOP-MULTI is an implicit coarse-grained model with a two-bead resolution for each amino acid except for glycine (represented by one bead), similar to previously described SOP-SC and SOP-IDP models. The model accommodates a spherical backbone (*B*) bead at the *C*_*α*_ position and a spherical side chain (*S*) bead at the centre of mass of the side chain atoms. SOP-MULTI contains force-field terms from both SOP-SC and SOP-IDP models, which include bonded (*E*_*bond*_), excluded volume (*E*_*exc*_), electrostatic(*E*_*electro*_) and Lennard-Jones potentials (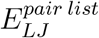 and 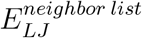).

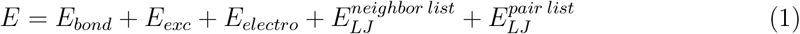

The critical addition to SOP-MULTI is the introduction of IDR-Folded domain interactions. This is addressed using hybrid beads for modeling folded domains. The beads belonging to the same folded domain use SOP-SC parameters for their interaction while they interact with other folded domains and IDRs using SOP-IDP parameters. The interaction between beads belonging to the same folded domain is defined using 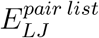. This potential is evaluated over a static pair list that is provided as input for the simulation. This pair list contains all the pairs of beads within a radius of *R*_*c*_ that belong to the same domain and are defined by SOP-SC parameters. On the other hand, 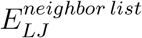 runs over a dynamically changing list of pairs and acts between all the pairs of beads belonging to either different domains or IDR and are within the cutoff provided to the potential. This potential uses a neighbour list generated by the molecular dynamics engine and uses parameters from SOP-IDP. Table 1 provides an illustrative picture of these interactions in an MDP with two well-folded domains (D1 and D2) and IDRs.

**Table 1:** Iteractions between beads belonging to IDRs, same domain, different domains and between IDR and folded domains.

|  | IDR-IDR | IDR-D1 | IDR-D2 | D1-D2 | D1-D1 | D2-D2 |
| --- | --- | --- | --- | --- | --- | --- |
| Bond | ✓ | ✓ | ✓ | ✓ | ✓ | ✓ |
| Excluded volume | ✓ | ✓ | ✓ | ✓ | ✓ | ✓ |
| Electrostatic | ✓ | ✓ | ✓ | ✓ | ✗ | ✗ |
| LJ : Neighbor list | ✓ | ✓ | ✓ | ✓ | ✗ | ✗ |
| LJ : Pair list | ✗ | ✗ | ✗ | ✗ | ✓ | ✓ |

The description of each of the terms in equation1 is as follows.

#### FENE Potential for bonded interaction (*E*_*bond*_)

The bonds among the backbone and sidechain beads (*B − B* and *B − S*) were modelled using finitely extensible nonlinear elastic (FENE) potential.^58^ This potential is active between all the connected beads of the protein.

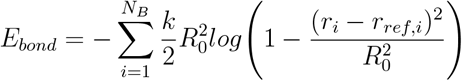

where *k* is the energy scaling factor, *R*_0_ is the tolerance displacement and *r*_*ref,i*_ is the reference bond distance between the connected beads of ‘*i*’th bond. The summation runs over all the *N*_*B*_ bonds of the bond list.

#### Excluded volume Potential (*E*_*exc*_)

Excluded volume is a purely repulsive potential that maintains the the volume of exclusion for each bead with defined radii(*σ*_*i*_) using the following function and is active between all pairs connected by 1-3 interaction.

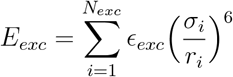

where *ϵ*_*exc*_ is the energy scaling factor of excluded volume potential, *σ*_*i*_ is the sum of the radii of the two beads involved in the interaction.

#### Electrostatic Potential (*E*_*electro*_)

The electrostatic potential is defined by a screened Coulomb potential between all the pairs of charged beads within the provided cutoff. It uses a neighbour list based approach to determine the pairs of interacting partners. This potential does not act on pairs belonging to the same folded domain. It is also inactive between pairs separated by less than 1-3 interactions.

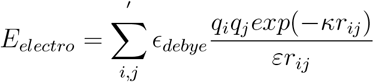

where *ϵ*_*debye*_ is energy scaling factor of the potential, *q*_*i*_ and *q*_*j*_ are the charges on beads *i* and *j, κ* is the inverse of Debye length and *ε* is the dielectric constant. *κ* and *ε* are a function of temperature and salt concentration. ^27,28^

#### Lennard-Jones Potential - Neighbour list based 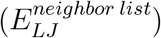

This Lennard-Jones potential acts on all pairs that are dynamically listed by the neighbour list of the molecular dynamics engine and is inactive among bead pairs belonging to the same domain.

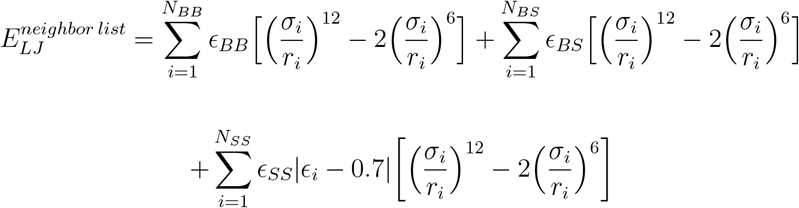

Where *ϵ*_*BB*_, *ϵ*_*BS*_ and *ϵ*_*SS*_ are scaling factors for Backbone-Backbone, Backbone-Sidechain, Sidechain-Sidechain interactions respectively. *ϵ*_*i*_ is the parameter that is a function of bead types involved in the pair and is adopted from Betancourt-Thirumalai statistical potential. ^59^

#### Lennard-Jones Potential - Pair list based 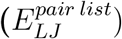

This potential defines the interaction among the beads of individual folded domains using a predefined list of interacting beads, which is provided as input for the simulation.

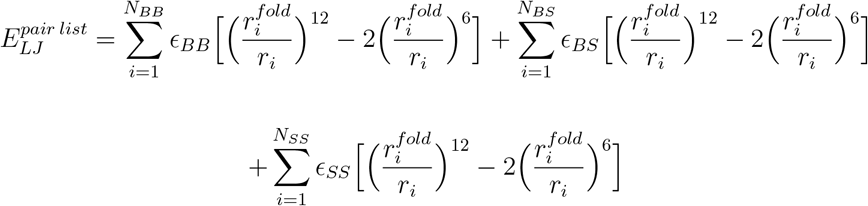

### 2.2 Architecture of SOP-MULTI

The architecture of SOP-MULTI (see Fig. 1) consists of two components: input files setup using Python scripts and the core C++ LAMMPS addon code. The input files setup (see Blue block in fig.1) includes generating a LAMMPS data file, a list of beads belonging to well-folded domains (for MDP only), and a forcefield file. The core add on code of SOP-MULTI user-package includes the features shown in the right block of fig. 1. A complete, detailed manual describing these features and usage was provided in the supporting information 2 (SI-2).

**Figure 1:**
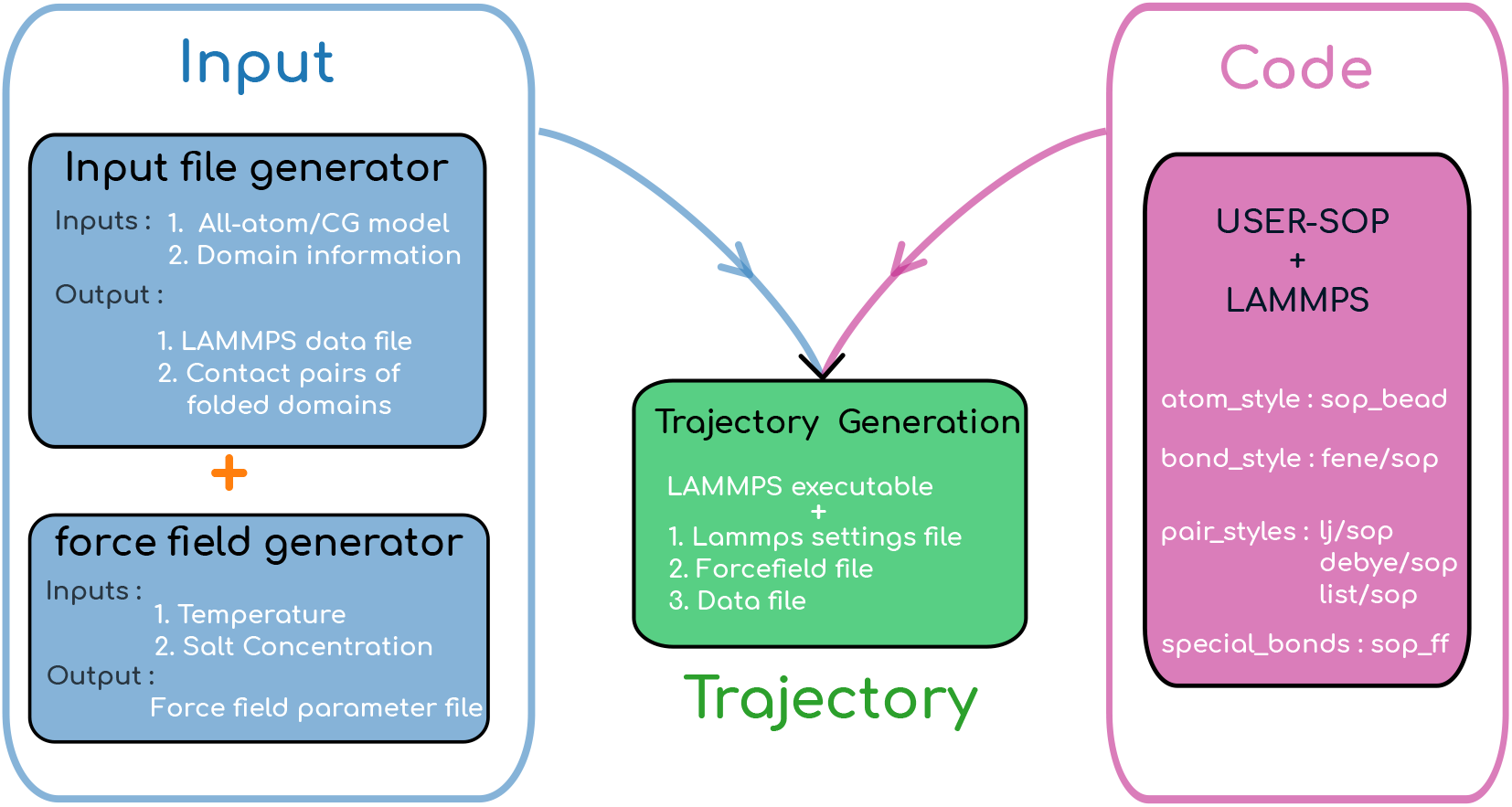
Architecture of SOP-MULTI package

### 2.3 Simulation Methodology

The initial all-atom resolution structures were generated by using either Alphafold’s colab notebook^60^ or modeling of structures using existing crystallographic structures of the folded domains. With these initial coordinates in hand, we developed an in-house Python code to generate an SOP resolution model of the given all-atom model and further generate LAMMPS-compatible input files. The trajectories were generated using LAMMPS patched with SOP-MULTI code. The trajectories were generated using Langevin dynamics with a timestep of 10 *fs*. Each of the systems was energy minimized using a conjugate gradient algorithm before proceeding to equilibration and production runs. Each of the systems was run for an order of 1 *−* 5 *×* 10^9^ steps. The auto-correlation function of the radius of gyration was monitored to ensure convergence of the simulated system. The initial equilibration phase of the trajectory was discarded before evaluating the polymer properties of the IDP/MDP.

All the LAMMPS settings can be found in the files deposited at the following link. https://github.com/codesrivastavalab/SOP-MULTI

### 2.4 Analysis

#### Contact Map

The contact map is generated by evaluating the average distance observed over the trajectory between all the possible pairs of backbone(C_*α*_) beads. These distances are visualized in the form of a heatmap. The intensity of colour represents the distance. In this article, the shorter distances are indicated by dark blue, which gradates into white for longer distances. The contact map allows us to check the intactness of folded domains and inter-domain interactions.

#### Scattering intensities calculation

The Scattering intensities of a given SOP-MULTI protein trajectory can be calculated using two methods. The first method uses the Debye formulation described by the equation below to evaluate the scattering intensity directly from the coarse-grained trajectory of the protein.

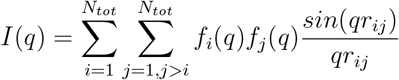

where *q* is the scattering vector, and *f* is the form factor of individual beads of the model. The form factors of individual beads were adopted from previous work. ^61^ A reproduction of the form factors of individual form factors can be found in Fig. S 4 of SI-1. In the second method, the CG trajectory is back-mapped to all-atom resolution using a reconstruction tool(PULCHRA^62^), which is then used to calculate scattering intensity using CRYSOL^63^ that uses a spherical harmonics-based approach. ^64^

#### Pair distance distribution function

Pair distance distribution function/curve *P* (*r*) is essentially a histogram of all the atom-atom pair distances (*r*) observed in a given protein by the beam during a small angle scattering experiment. It is obtained by evaluating the inverse transform of the one dimensional *I*(*q*) (Intensity as a function of scattering vector *q*). Simulation trajectories are obtained by histogramming all the pair distances observed in a given snapshot.

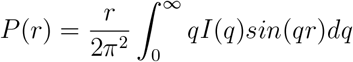

This function is evaluated from the experimental one-dimensional small-angle scattering data using SAS processing tools^65^ GNOM.^66^ On the other hand, the PDDF is evaluated from the simulations by simply evaluating all the pairwise distances between all the atoms of the polymer and histogramming these distances.

#### Shape characteristics of single molecule

The mean properties of the protein polymers are captured by the following properties, which can be evaluated from simulations. The gyration radius of a given protein snapshot was evaluated using the following relation with the appropriate mass of the individual CG beads of residues.^28^ COM is the centre of mass of the protein, while *m* and *r* represent the bead’s mass and position vector, respectively.

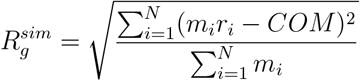

#### Hydrodynamic radius

The hydrodynamic radii of the proteins were evaluated using the following relation.

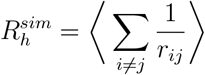

#### Asphericity and Shape parameters

The asphericity (Δ) and the shape parameter (*S*), which are both derived from the inertia tensor, define deviation from the spherical shape. Previous studies have shown the applicability of these parameters to RNA^67,68^ and protein^69^ molecules in characterizing their shape. Asphericity and Shape parameter are defined as follows.

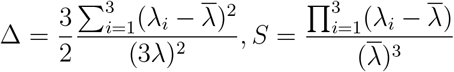

where *λ*_*i*_ are the eigenvalues of the gyration tensor and 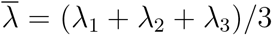 The gyration tensor^70^ is given by the following matrix

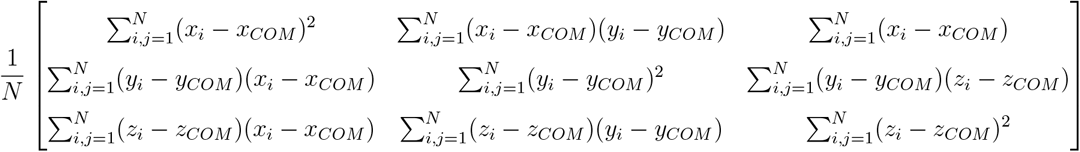

Asphericity lies in the range 0 *≤* Δ *≤* 1, where 0 represents a spherical geometry, and 1 represents a linear rod-like geometry. The shape parameter has the bound of *−*1*/*4 *≤ S ≤* 2, where negative and positive values of the shape parameter indicate oblate and prolate shapes, respectively.

#### Scaling exponent

The Scaling exponent was calculated using two methods. The first one uses the mean radius of gyration observed with different IDPs of varying lengths.

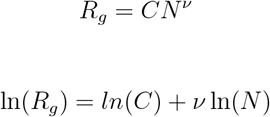

In the second method, we evaluate the distribution of scaling exponents using multiple snapshots of the trajectory of the protein of interest. The average distance between two residues separated by varying numbers of residues was used to evaluate the exponent by fitting it to the following exponent.

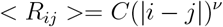

An example of such fitting for histatin5 IDP, along with the limits for compact globule, good solvent and flory limit, is shown in Figure 2 of SI. The scaling exponent code was developed by applying the needed modifications to the source code of SOURSOP**?**

**Figure 2:**
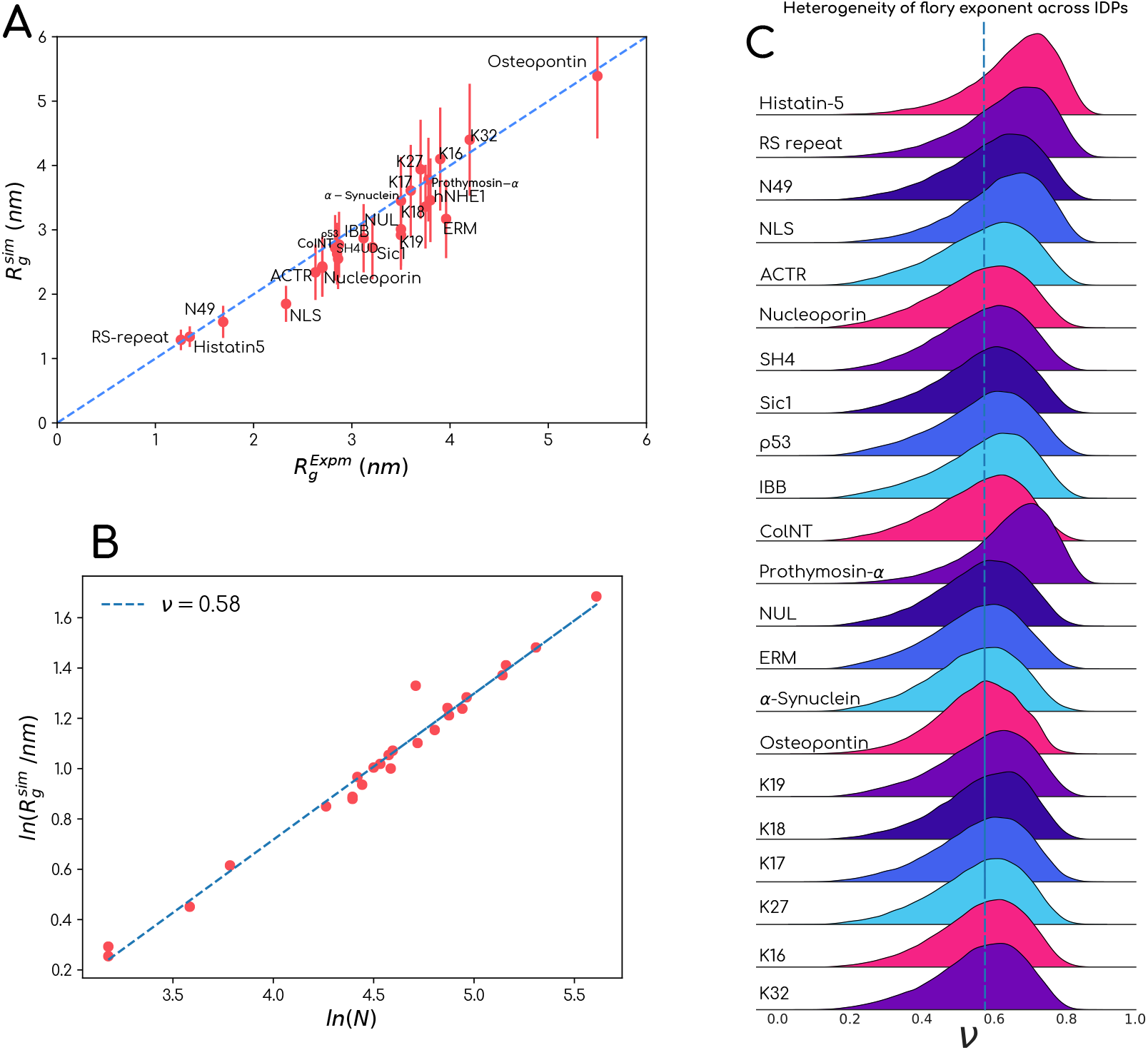
A. Comparison of the experimentally observed radius of gyration with that obtained from SOP-MULTI trajectories of corresponding IDPs B. Flory exponent derived using trajectories all the IDPs using SOP-MULTI model. C. Distribution of flory exponents observed in the trajectories of individual IDPs.

#### Root Mean Square Fluctuations

The Root Mean Square Fluctuations (RMSF) in CG simulations are calculated using the RMSF module of MTraj package.^71^ The RMSF is calculated using the variance in positions *x*_*i*_ of backbone beads of individual residues from their mean positions ⟨*x*_*i*_ ⟩.

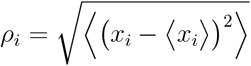

## 3 Results and discussion

### 3.1 Intrinsically Disordered Proteins

### 3.1.1 SOP-MULTI generates IDP ensembles consistent with SOP-IDP

The SOP-MULTI code was tested on 22 IDPs of varying lengths chosen from previous SOP-IDP studies.^27,28^ A table of the simulated IDP systems and their experimental *R*_*g*_, along with that observed with SOP-MULTI, can be found in Table 2. The observed 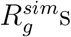 are in good agreement with previous studies performed in OPENMM^72^ environment. A com-parison of the experimental 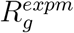 and simulation 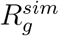 can be observed in Fig. 2A. We noticed that there is good agreement between both of them. We further checked the Flory scaling exponent of 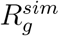 with the increasing length of IDP using trajectories of all the 22 IDP simulations and obtained ν = 0.58 (Fig. 2B), which indicates that these IDPs behave as polymers in a good solvent. With all the trajectories in hand, we also looked at the distribution of Flory exponents of individual conformations of a given IDP, and we observed that every IDP dynamically samples a distribution of exponent values with mean lying in the good solvent regime (Fig. 2C). This shows the heterogeneity in the IDP(R) ensemble that arises due to variation in sequences. Heterogeneity in other polymer properties, such as asphericity and shape parameters, was also observed (see Figure 1 in SI-1). Such observations using conformational ensembles of 28058 IDP(R)s found across human proteome were reported in a recent study.^9^ Put together, our simulations show that SOP-MULTI works for simulating pure IDPs, and the results are as good as the source SOP-IDP model.

**Table 2:** Intrinsically disordered protein systems tested with SOP-MULTI

|  | Protein | Length | Temperature (°C) | Salt Conc. (mM) |
| --- | --- | --- | --- | --- |
| 1 | Histatin5<sup>23</sup> | 24 | 25 | 150 |
| 2 | RS-repeat<sup>73</sup> | 24 | 25 | 150 |
| 3 | N49<sup>74</sup> | 36 | 25 | 150 |
| 4 | NLS<sup>74</sup> | 44 | 25 | 150 |
| 5 | ACTR<sup>75</sup> | 71 | 5 | 500 |
| 6 | Nucleoporin<sup>74</sup> | 81 | 25 | 150 |
| 7 | SH4UD<sup>76</sup> | 85 | 25 | 150 |
| 8 | Sic1<sup>77</sup> | 90 | 25 | 150 |
| 9 | p53<sup>78</sup> | 93 | 25 | 150 |
| 10 | IBB<sup>74</sup> | 97 | 25 | 150 |
| 11 | CoINT<sup>79</sup> | 98 | 4 | 400 |
| 12 | Prothymosin-α<sup>80</sup> | 111 | 25 | 150 |
| 13 | NUL<sup>74</sup> | 112 | 25 | 150 |
| 14 | ERM<sup>81</sup> | 122 | 25 | 150 |
| 15 | α-Synuclein<sup>82</sup> | 140 | 25 | 200 |
| 16 | Osteopontin<sup>83</sup> | 273 | 25 | 80 |
| 17 | K19<sup>84</sup> | 99 | 15 | 150 |
| 18 | K18<sup>84</sup> | 130 | 15 | 150 |
| 19 | K17<sup>84</sup> | 143 | 15 | 150 |
| 20 | K27<sup>84</sup> | 171 | 15 | 150 |
| 21 | K16<sup>84</sup> | 174 | 15 | 150 |
| 22 | K32<sup>84</sup> | 202 | 15 | 150 |

### 3.2 Multidomain Proteins

We simulated multiple full-length MDPs using the SOP-MULTI package. A list of MDPs simulated with SOP-MULTI as part of this work can be found in Table 3. The domain architecture and a snapshot of the expanded form of the SOP-MULTI model of each of these MDPs are reported in subfigures A and B respectively of Figures 3-11. The proteins vary in length and domain architecture. Proteins with only one folded region flanked by IDRs were also considered as MDPs in our study.

**Table 3:** Multidomain proteins systems simulated with SOP-MULTI

|  | Protein | Length | no. IDRs | Domains | Temp (°C) | Salt Conc. (mM) |
| --- | --- | --- | --- | --- | --- | --- |
| 1 7 | Ubiquitin<sub>2</sub><sup>85</sup> | 162 | 3 | 2 | 20 | 300 |
| 2 7 | Ubiquitin<sub>3</sub><sup>85</sup> | 228 | 3 | 3 | 20 | 300 |
| 3 10 | hGHR-ECD<sup>86</sup> | 244 | 2 | 1 | 25 | 150 |
| 4 6 | TIA-1<sup>87,88</sup> | 275 | 4 | 3 | 20 | 100 |
| 5 7 | Ubiquitin<sub>4</sub><sup>85</sup> | 304 | 4 | 4 | 20 | 300 |
| 6 9 | hnNRP A1<sup>18</sup> | 314 | 3 | 2 | 20 | 150 |
| 7 3 | TDP-43<sup>89</sup> | 414 | 4 | 4 | 27 | 100 |
| 8 4 | HIV-GAG<sup>90</sup> | 432 | 4 | 5 | 25 | 100 |
| 9 8 | G3BP1<sup>91</sup> | 466 | 2 | 2 | 25 | 1 |
| 10 8 | G3BP1<sup>91</sup> | 466 | 2 | 2 | 25 | 10 |
| 11 8 | G3BP1<sup>91</sup> | 466 | 2 | 2 | 25 | 100 |
| 12 8 | G3BP1<sup>91</sup> | 466 | 2 | 2 | 25 | 1000 |
| 13 11 | Human FUS | 526 | 3 | 2 | 25 | 150 |

**Figure 3:**
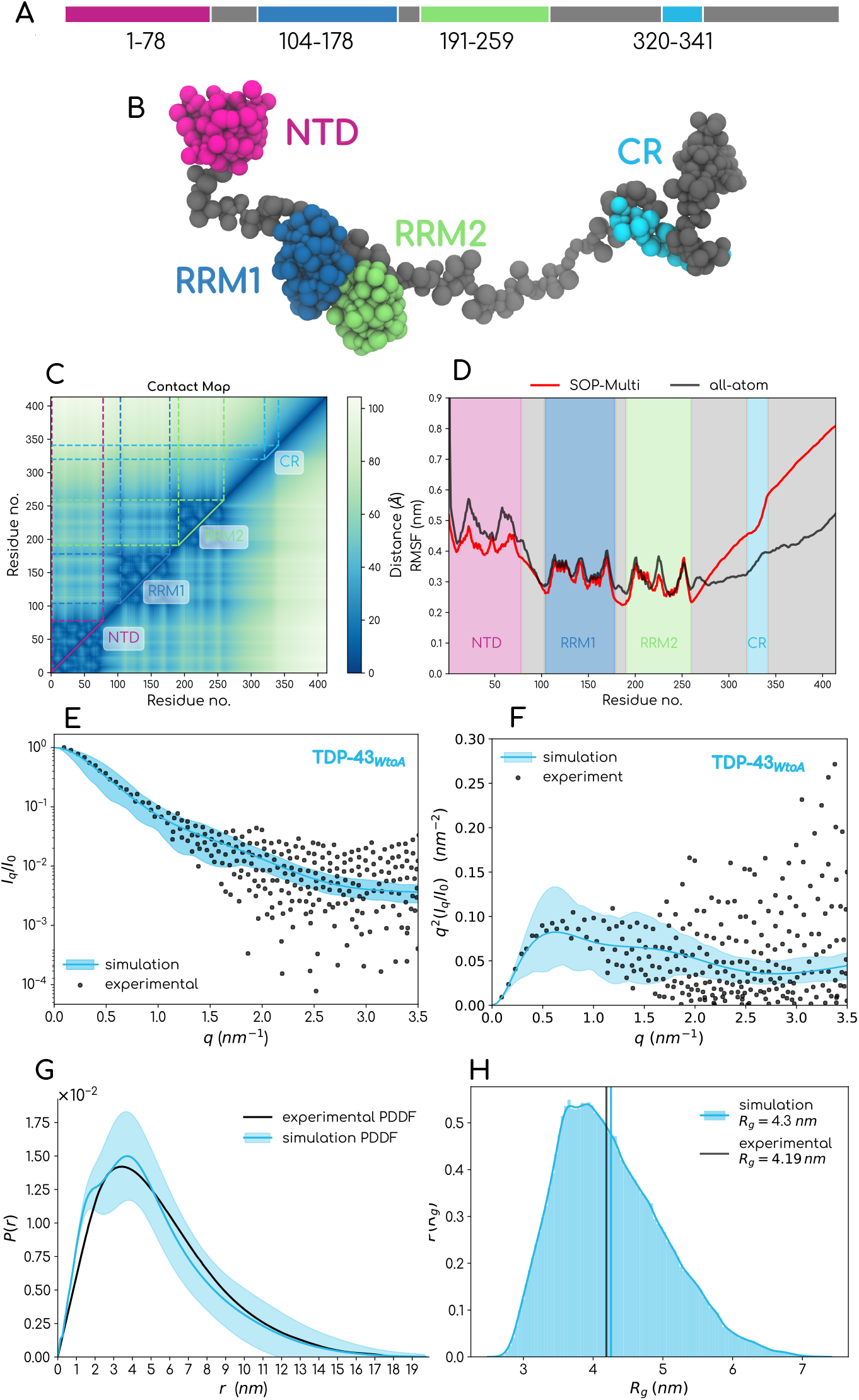
TDP-43: A. Domain architecture B. SOP model C. Contact map D. Comparison of scattering Intensity E. Kratky plot F. Pair distance distribution function G. Radius of Gyration

#### 3.2.1 SOP-MULTI maintains stable folded domains in full length MDPs

As mentioned in the previous section, the interactions among the beads of individual folded domains are maintained by a static neighbour list of Lennard-Jones pairs. We checked the stability of the folded regions by looking at the contact maps of these proteins and checking if the intra-domain contacts are maintained. These maps were reported in subfigure C of Figures 3-11. Dotted lines and solid lines across the diagonals of these contact maps indicate the folded domains. As evident from these contact maps, the folded domains remain intact through the trajectory of these MDPs, inferring that the three-dimensional structure of folded domains is preserved. These maps also aid in quantifying/visualizing the inter-domain, intra-domain and IDR-domain interactions.

#### 3.2.2 SOP-MULTI is capable of generating scattering data compatible conformation ensembles of full length MDPs

TDP-43 is a multidomain protein with four folded regions, two long IDRs and two linker stretches(also modelled as IDRs). The protein starts with an N-terminal domain (NTD) followed by two RNA recognition motifs (RRM1 and RRM2) connected by two linker regions. The RRM2 is followed by a conserved region(CR) flanked by two IDRs. Accumulation of aggregates of this MDP in the central nervous system is a typical signature observed in many neurodegenerative diseases.^13^ To probe biophysical insights into this protein, purification of aggregation-free full-length TDP-43 was facilitated by mutating all the six tryptophans present in this MDP to alanine.^89^ We use the scattering data from this study to see the performance of the SOP model of TDP-43_*W toA*_ (Fig. 3B) in capturing the global single molecule properties. We simulated the trajectory of TDP-43_*W toA*_ and evaluated the theoretical scattering intensity from the trajectory. We then compared the evaluated scattering intensity with experimental data. We found a consistent match across the entire reciprocal regime with raw and Kratky-transformed intensity data, see Fig. 3E and F. This match was also observed when the scattering intensities were translated to real space PDDF function (Fig. 3G). The mean radius of gyration(*R*_*g*_ = 4.3 *nm*) observed in the simulated SOP model of TDP-43 is quite close to the (*R*_*g*_ = 4.19 *nm*) derived from scattering data (Fig. 3H). These indicate that SOP-MULTI could capture a scattering realistic ensemble of TDP-43.

HIV1-GAG is another MDP with similar architecture as that of TDP-43 (Fig. 4A) and is of active interest for its role in assembly of HIV-1 virus. ^92^ It is a polyprotein with three folded domains matrix (MA), capsid (CA) and nucleocapsid (NC) and three shorter peptides SP1, SP2 and p6.^93^ A structual exploration of a solution state single molecule properties was made possible with scattering methods^94^ and generation of mutant HIV-GAG_*W M*_, (lacking P6, tryptophan-316 and methionine-317 (184 and 185 of CA) mutated to alanine)) that retains the properties of wild type protein. The SOP-MULTI model could capture scattering data consistent ensemble as evident from intensity plot (E), Kratky plot(F) and PDDF plot (G) in Fig. 4. The mean *R*_*g*_ observed with SOP-MULTI is close to *R*_*g*_ derived from Guinier region of scattering data.

**Figure 4:**
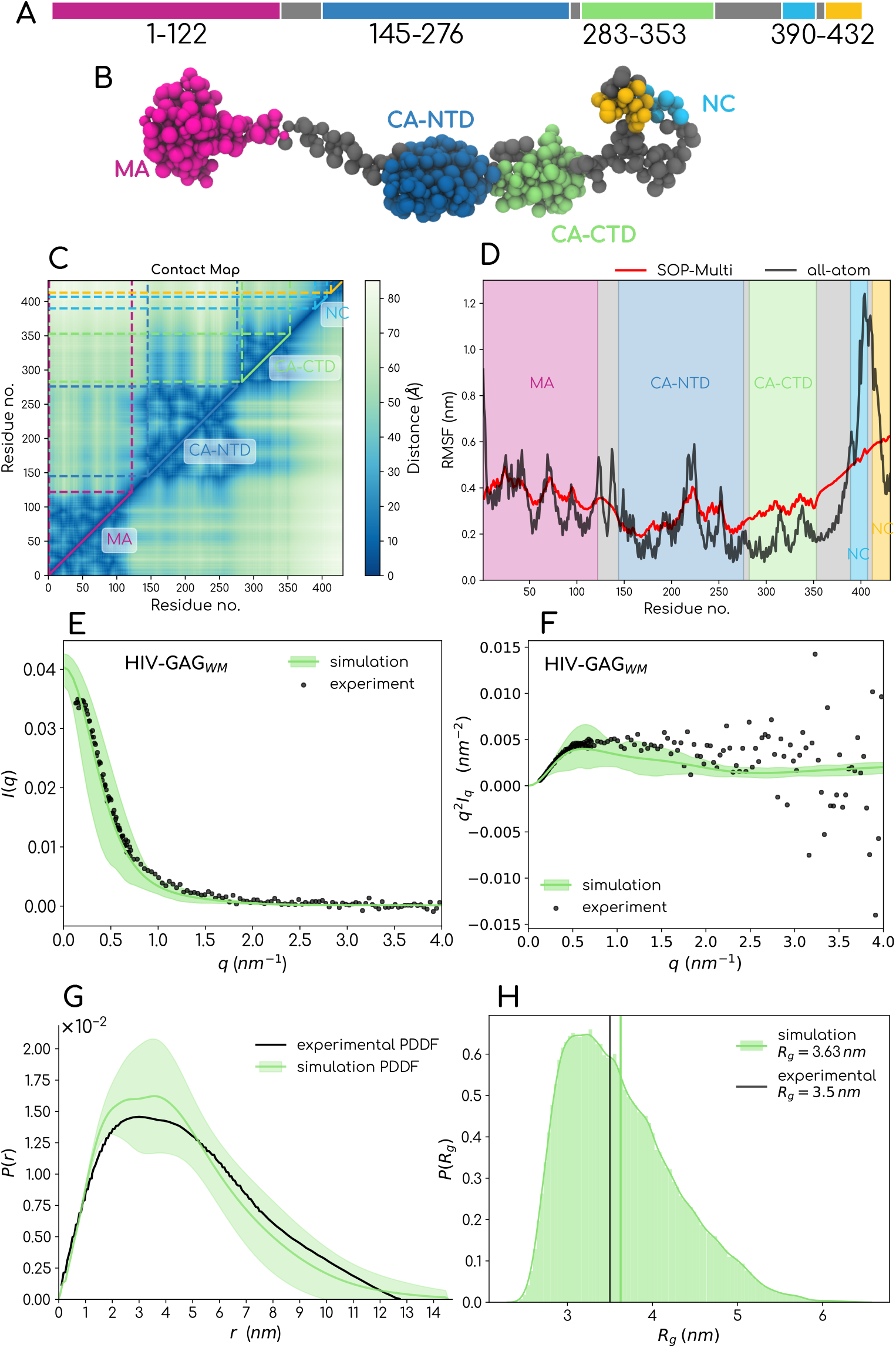
HIV-GAG: A. Domain architecture B. SOP model C. Contact map D. RMSF comparison with all-atom model E. Comparison of scattering Intensity F. Kratky plot G. Pair distance distribution function H. Radius19of Gyration

#### 3.2.3 SOP-MULTI captures trends in all-atom fluctuations

SOP-MULTI models individual domains using the SOP-SC model that preserves the native contact information. The rigorous optimization of energy scales from the previous works allowed us to capture the trend in all-atom fluctuations, too. We used all-atom trajectories of TDP-43^95^ and HIV1-GAG^*to be published*^ to observe the fluctuations in the backbone of these MDPs. We then compared these fluctuations with those evaluated from SOP-MULTI trajectories (shown in red in Figures. 3D and HIV-GAG4D). We observed that the trend in fluctuations of folded domains is well captured with SOP-MULTI in comparison with all-atom simulations(shown in black in Figures. 3D and 4D). However the IDRs in these MDPs have higher fluctuations in SOP-MULTI. This can be attributed to the smoother energy landscapes of these proteins in coarse-grained space, which allows for a much faster sampling of the conformational landscape than the rugged energy landscape. This can be observed in the case of TDP-43 Fig.3D and HIV-GAG4D where a comparison of root mean square fluctuations was performed between SOP-MULTI and all-atom simulations.

#### 3.2.4 All-atom reconstruction of SOP-MULTI trajectories recapitulates scattering data over a wider range of reciprocal space

To check the experimentally realistic nature of trajectories generated using the SOP-MULTI model, we evaluated the scattering intensities using CG form factors reported previously^61^ of TDP-43 and TIA-1 MDPs. We notice (see Fig. 5) that the evaluated scattering profiles match well in the lower *q* regime (0.0 *−* 1.5 *nm*^*−*1^) but deviate from the experimental data at higher *q* regime (*>* 2.0 *nm*^*−*1^), which indicates a relatively collapsed ensemble compared to experimental data. We went on to investigate this further by back-mapping the trajectory to all-atom resolution and then seeing if the deviation was reduced. We do notice an improvement in the match with the reconstructed trajectory, indicating the need for a better forward model for calculating scattering intensities from CG trajectories. The possible reason for this is the training dataset considered for optimizing these CG form factors is populated with folded proteins. The form factors were reported to be successfully applied to IDPs. However, they failed to capture the high *q* regime for multidomain proteins. This is a key area that needs to be addressed to guide efforts in developing forcefields consistent with scattering data in the right direction.

**Figure 5:**
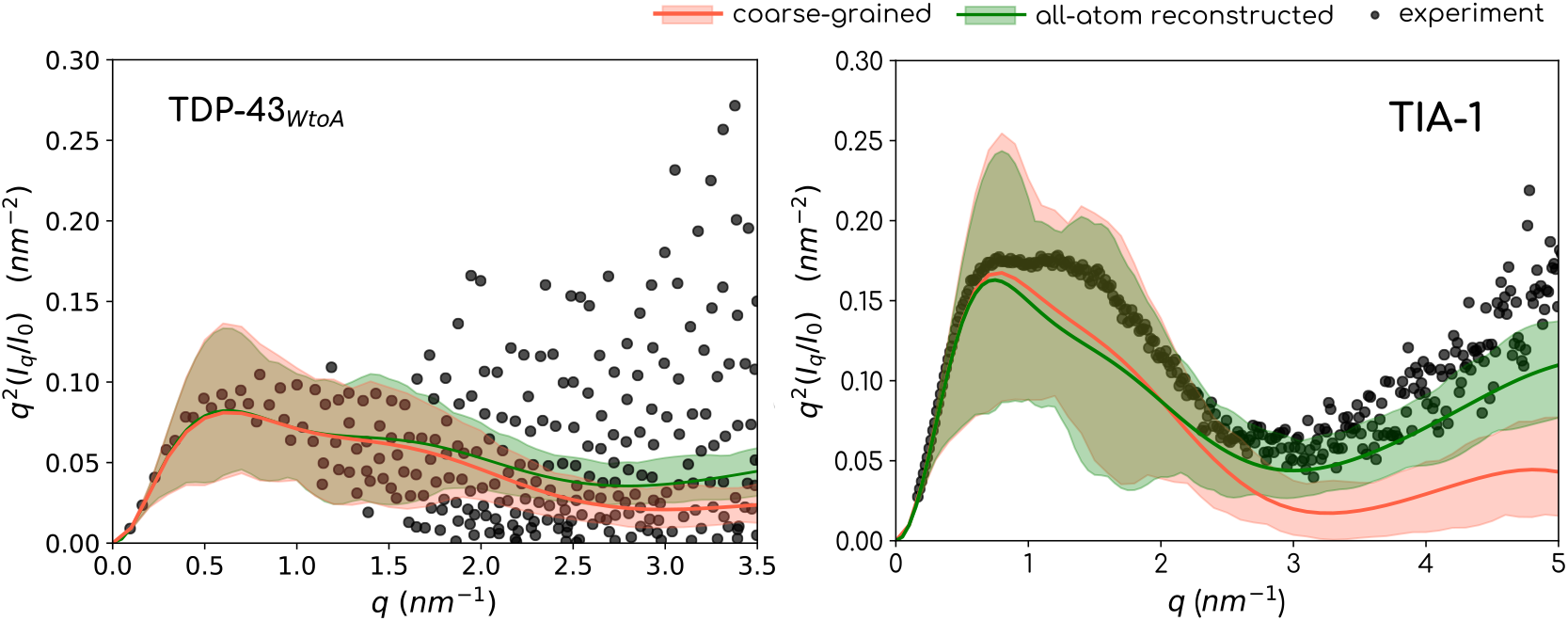
Kratky representation of simulation scattering intensities of TIA-1(Right) and TDP-43(Left) SOP-MULTI ensemble with experimental data. Red : Using CG trajectory and CG form factors for evaluation of scattering intensities. Green : Using all-atom reconstructed trajectory and atomic form factors

#### 3.2.5 SOP-MULTI captures ensembles comparable to optimized Martini simulations: Use cases as TIA-1 and Poly-Ubiquitin

TIA-1 is an MDP with three RNA recognition motifs connected by flexible linker regions (Fig. 6A). Thorough scattering studies have been performed using X-ray and neutron scattering along with computational modeling^87^ to probe the relative positions of individual domains. Martini model of this MDP was shown to sample compact conformations, which was then improved by increasing the strength of protein-water interactions.^88^ We generated the SOP-MULTI model of this MDP (see Fig. 6A) and found that the ensemble matches the experimental ensemble in both real(Fig. 6G) and reciprocal space(Fig. 6E, F). The inter domain distance between RRM1 and RRM3 domains(Fig. 6D) and the radius of gyration(Fig. 6H) match excellently with reported data from experiments and optimized Martini models.

**Figure 6:**
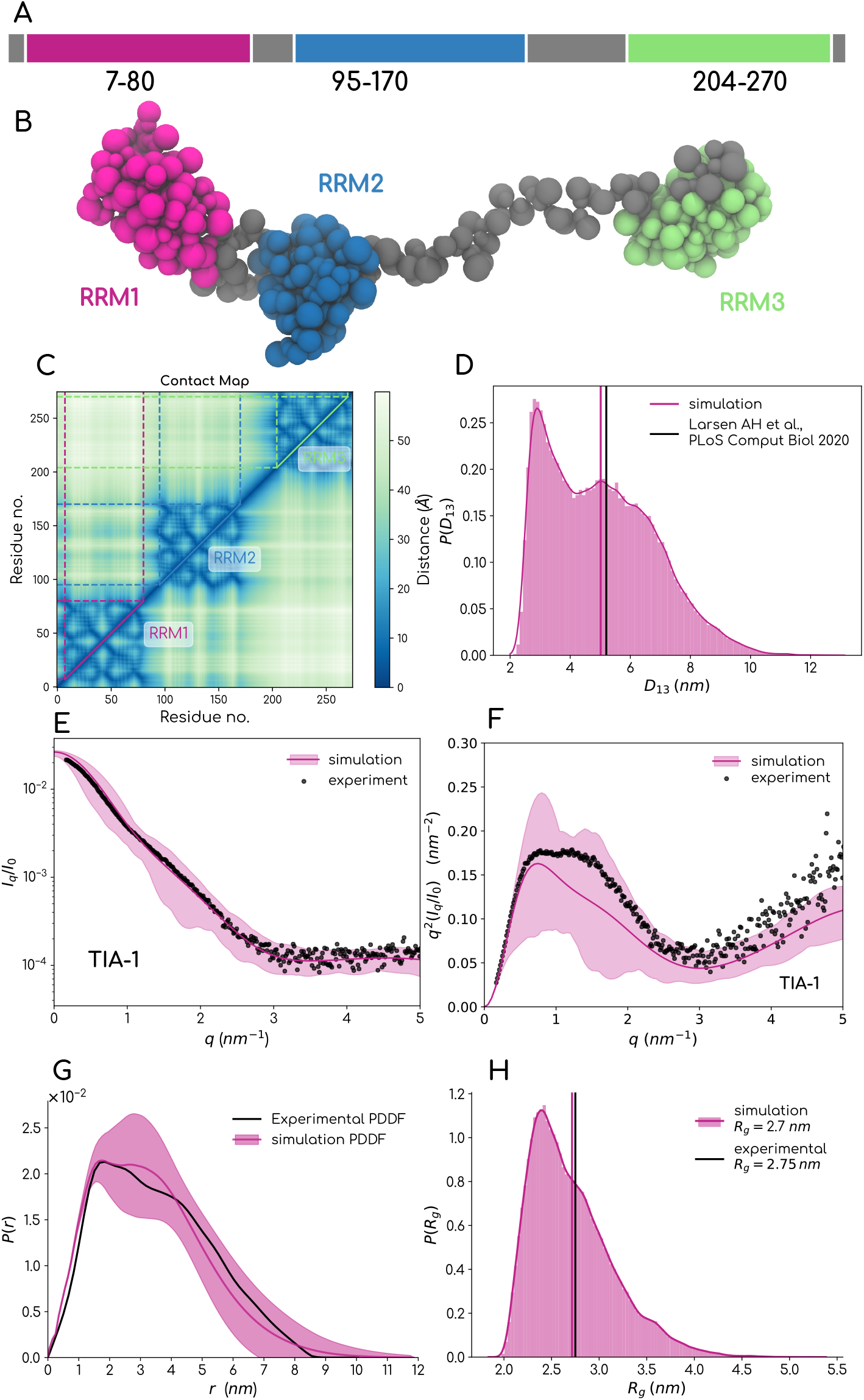
TIA-1: A. Domain architecture B. SOP model C. Contact map D. Comparison of scattering Intensity E. Kratky plot F. Pair distance distribution function G. Radius of Gyration

Ubiquitin, as the name says, is found ubiquitously across eukaryotic organisms^96^ and is a 76-residue-long, well-folded protein. The term ubiquitylation refers to adding one or more (polyubiquitin) of these to a target protein. The site of addition and length of polyubiquitin affects the target protein’s cellular localization and activity and alters its interactome. ^97^ Its well-known function is its ability to tag target proteins to be degraded by proteasomal machinery to ensure the recycling of proteins. After the first Ubiquitin binds to the target protein covalently, further ubiquitins link to one of its seven lysines or Methionine-1 (K6, K11, K27, K29, K33, K48, and K63 or M1). Polyubiquitins are thus an attractive model system for studying the biophysical characteristics of proteins obtained with a folded unit as its monomeric unit. An integrative approach using X-ray scattering and Martini simulations were taken up in a very recent study to understand the dynamics of linear polyubiquitin with two(Ub_2_), three (Ub_3_) and four ubiquitins(Ub_4_).^85^ The authors report data from three types of simulations, i.e. 1. Martini^98^ 2. Martini with tuned protein-water interactions^88^ 3. Metadynamic metainference^99,100^ by integrating experimental data. We compare the SOP-MULTI model of these poly ubiquitins with these observations, and we notice that the SOP-MULTI captures the mean *R*_*g*_ of the ensemble (refer fig. 7D)comparable to the optimized Martini simulations from the previous work. The Ub_4_ presents a challenge in modelling studies of both Martini and SOP-MULTI, as can be observed in Figs. 7D and G.

**Figure 7:**
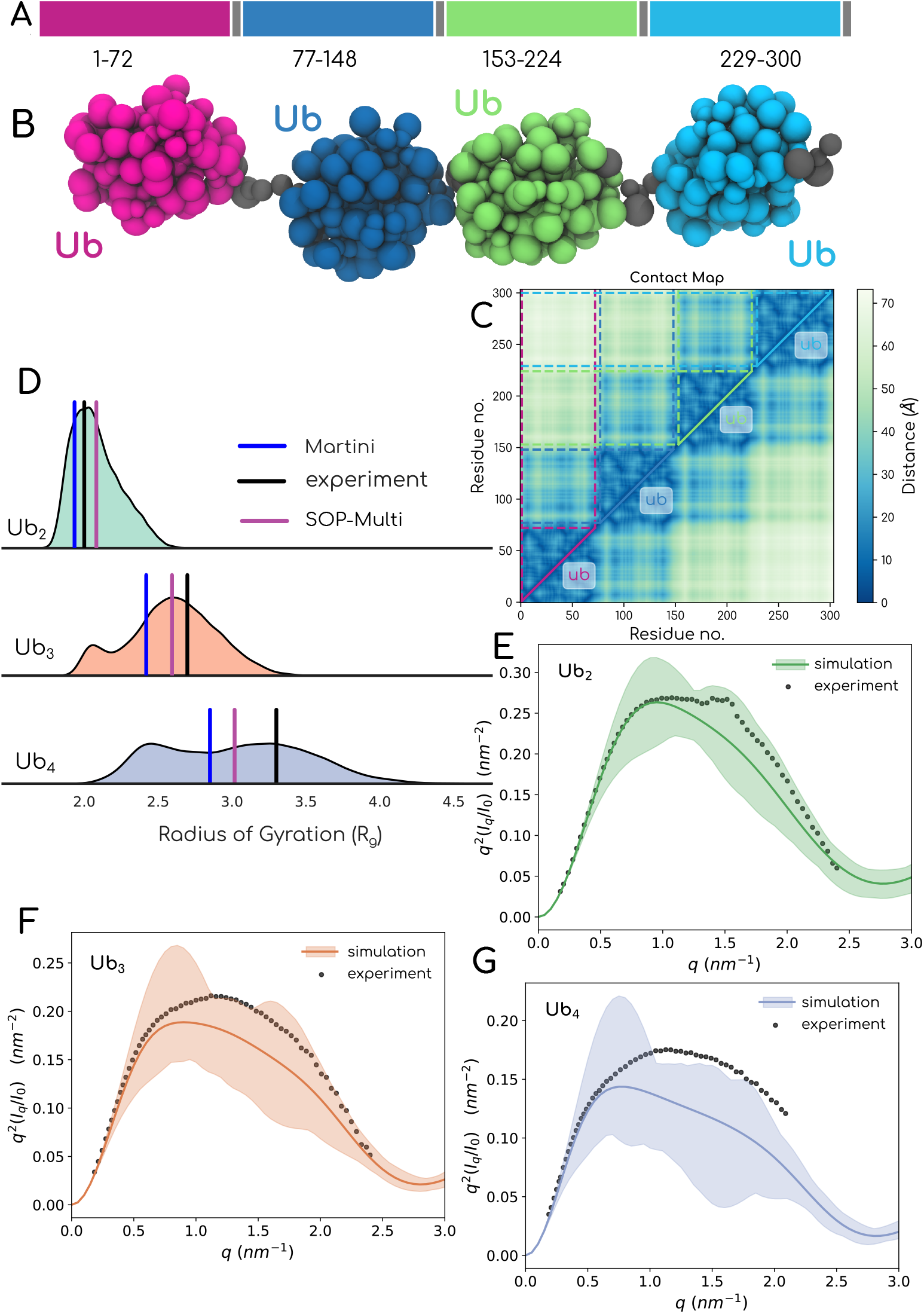
Polyubiquitin: A. Domain architecture of poly ubiquitins B. SOP model C. Contact map D. Comparison of *R*_*g*_ with experimental and Martini systems E-G. Kratky plot of Ub_2_, Ub_3_ and Ub_4_

#### 3.2.6 SOP-MULTI captures switching behaviour of G3BP1 with varying salt concentration

G3BP1, an RNA-binding protein, was shown to be one of the central players in the assembly of dynamic and reversible ribonucleoprotein stress granules in response to stress in eukaryotic cells. As part of understanding the interactome landscape of this protein, previous studies^91^ have observed a switch-like behaviour of this protein with a logarithmic increase in salt concentration. We modelled G3BP1 as MDP with two folded domains separated by a long IDR stretch of around 200 residues (Fig. 8A, B). The domain boundaries between folded and IDRs were determined using PAE and PLDDT deposited with the Alphafold^101,102^ model (see Fig. S3 in SI1). We simulated G3BP1 with increasing salt concentrations and found a similar switch-like transition of G3BP1’s ensemble from collapsed to expanded state (see intensity curves in Fig. 8D). A comparison of the calculated scattering intensity at low and high salt concentrations with experimental data in the form of dimensionless Kratky is reported in Figure 8D.

**Figure 8:**
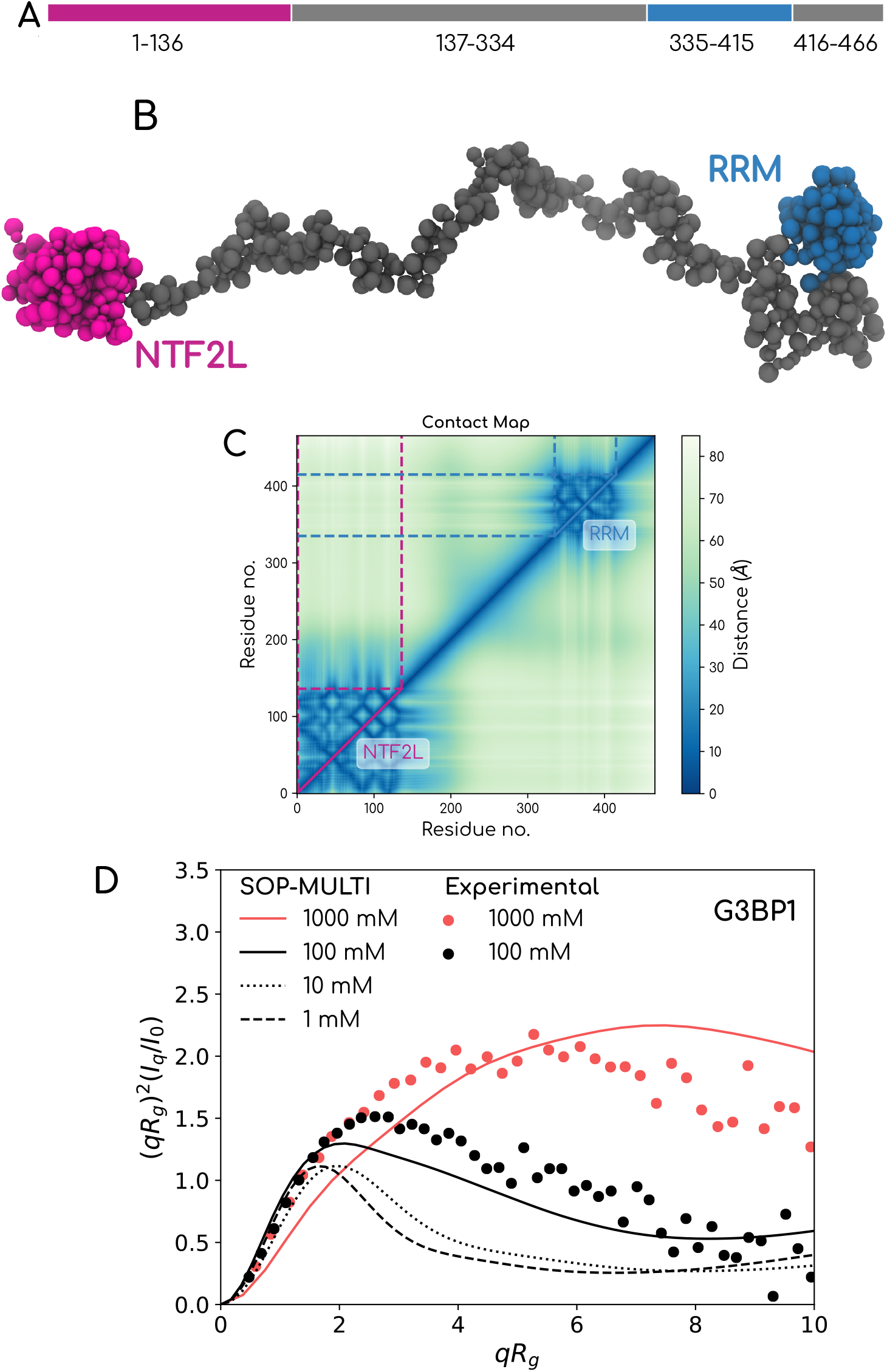
Switching Behaviour of G3BP1 from compact state to expanded state observed with change in salt concentration. Dimensionless Kratky representation of computed scattering intensities of G3BP1 observed at 1000 *mM* and 100 *mM* salt concentration overlaid with experimental data^91^

#### 3.2.7 SOP-MULTI captures conformational ensemble of folded protein with long IDR tails

MDPs are generally defined as proteins with more than one folded protein. However, we do have a repertoire of proteins with single-folded domains flanked by IDRs. We set forth to check if SOP-MULTI is capable of capturing the conformational ensemble of such proteins. We chose heterogeneous nuclear ribonucleoprotein (hnRNP A1) and human growth hormone receptor’s extracellular domain(hGHR-ECD) proteins. The domain architecture of both these proteins can be observed in figures 9A and 10A. The common feature is that both proteins contain one domain flanked on either end by IDRs. Defining domain boundaries is a crucial aspect of generating the SOP-MULTI model, as this determines how forces are described in the model. We used AlphaFold’s predicted aligned error(PAE)^1^ and predicted local distance difference test score (pLDDT)^103^ data to determine the number of domains and their boundaries of hnRNP A1(see Figure. 3a and 3b). PAE provides an estimate of the error in the position of each amino acid with respect to other residues, while pLDDT assigns confidence in the accuracy of prediction for that residue. A patch of low pLDDT score indicates a disordered region. Despite the typical nomenclature of the two domains of hnRNP A1 as RRM1 and RRM2, we consider both RRMs as single domain in our model. The other protein with a similar architecture is hGHR-ECD. Both these proteins have previous reports of scattering data, which allowed us to validate our model with proteins with such architecture. hGHR-ECD is well consistent with both scattering data in reciprocal space as well as real space PDDF and *R*_*g*_ data(see Figs. 10 D-G). However, in the case of hnRNP A1(9), we do notice that SOP-MULTI captures the mean *R*_*g*_, which has a relative error of *∼* 12% in comparison with the experimental value. However, the SOP-MULTI ensemble captures the entire scattering vector space accurately. We attribute the expanded ensemble of hnRNP A1 to the Low Complexity Region(LCR) present towards the C-terminal end of RRMs. The LCR was reported^104^ to be a special case of IDP, which is significantly collapsed compared with the vast majority of IDRome. The inability of SOP-IDP to capture this special case might be a possible reason for high *R*_*g*_ of the full-length hnRNP A1.

**Figure 9:**
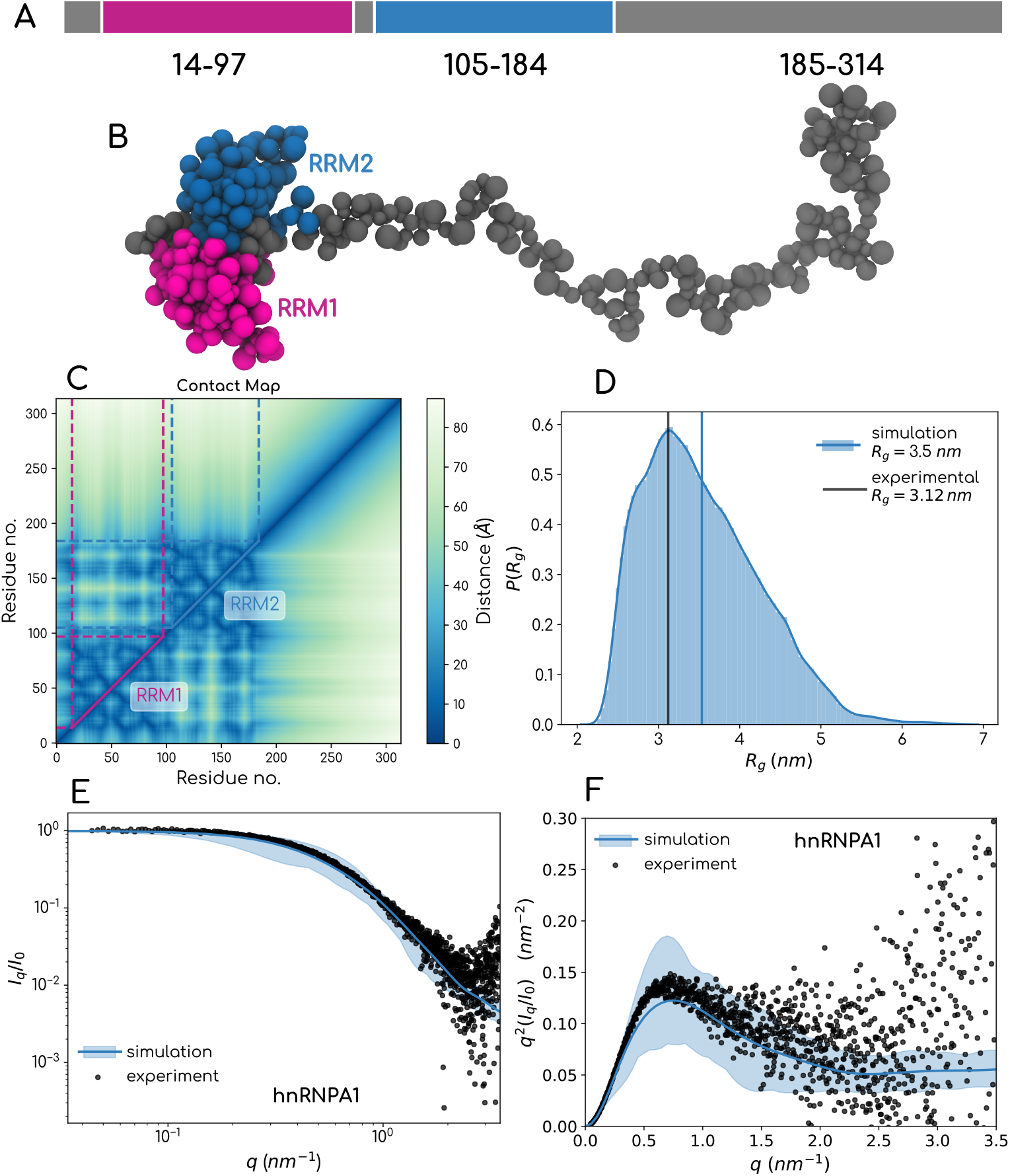
hnRNPA1: A. Domain architecture B. SOP model C. Comparison of scattering Intensity D. Kratky plot E. Radius of Gyration

**Figure 10:**
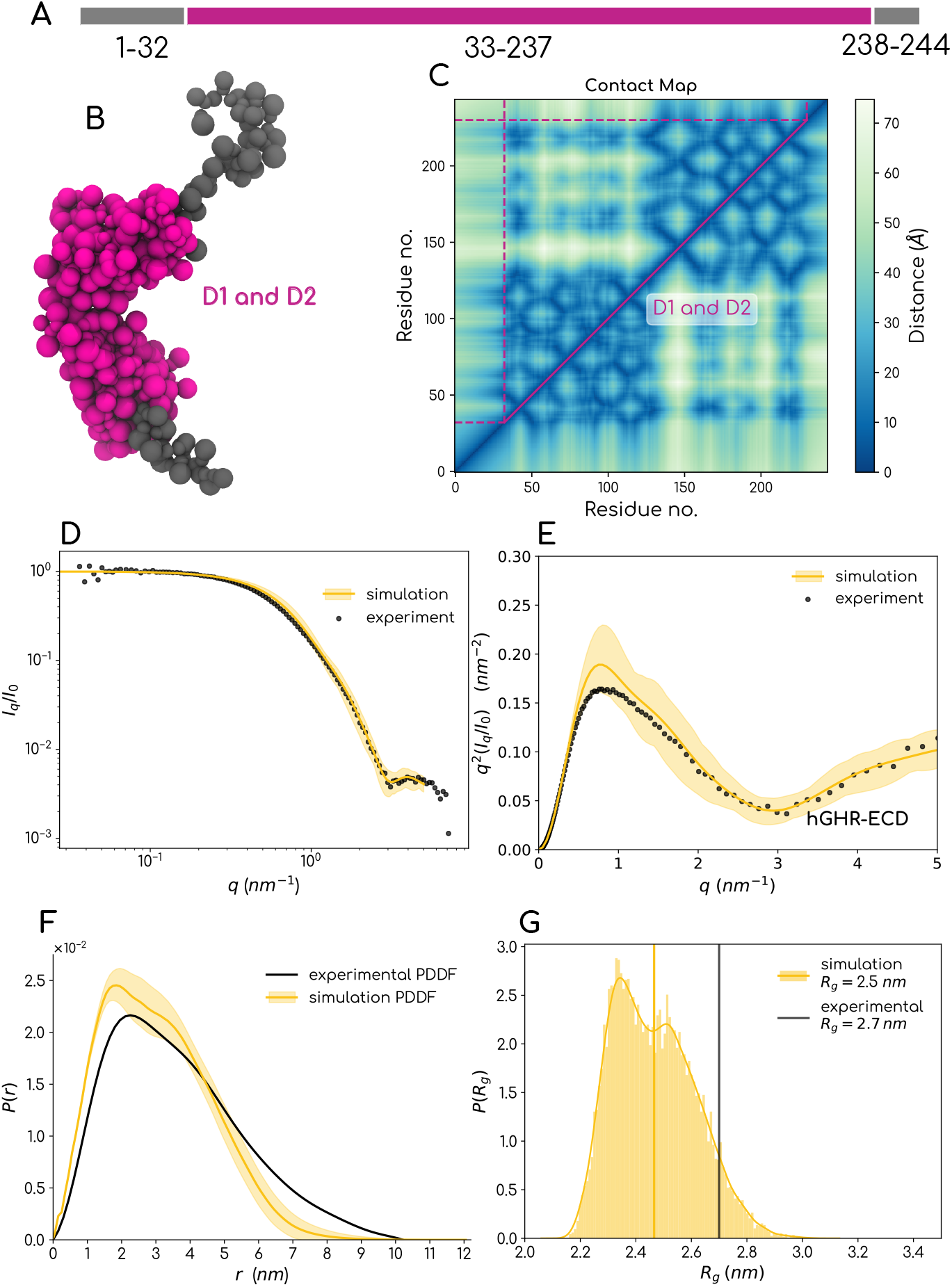
hGHR-ECD: A. Domain architecture B. SOP model C. Contact map D. Comparison of scattering Intensity E. Kratky plot F. Pair distance distribution function G. Radius of Gyration

#### 3.2.8 Prediction of single-molecule properties of Human-FUS

With SOP-MULTI as an ensemble generator, we predict the single-molecule properties of human-FUS MDP. FUS contains two folded domains, an RNA recognition motif and a zinc finger towards the C-terminal end (Fig. 11). We predict the *R*_*g*_ of human FUS to be in the range of 5 to 6 *nm* considering the mean and mode of the *R*_*g*_ distribution observed in the FUS simulation(see Fig. 11D). The presence of plateau-like behaviour in the medium to high *q* regime indicates the presence of higher populations of expanded conformations with an overall behaviour similar to that of an IDP. In essence, FUS behaves like an IDP despite the presence of two folded domains in the C-terminal region. Our predictions can be tested once an aggregation-free mutant of human FUS can be experimentally figured out.

**Figure 11:**
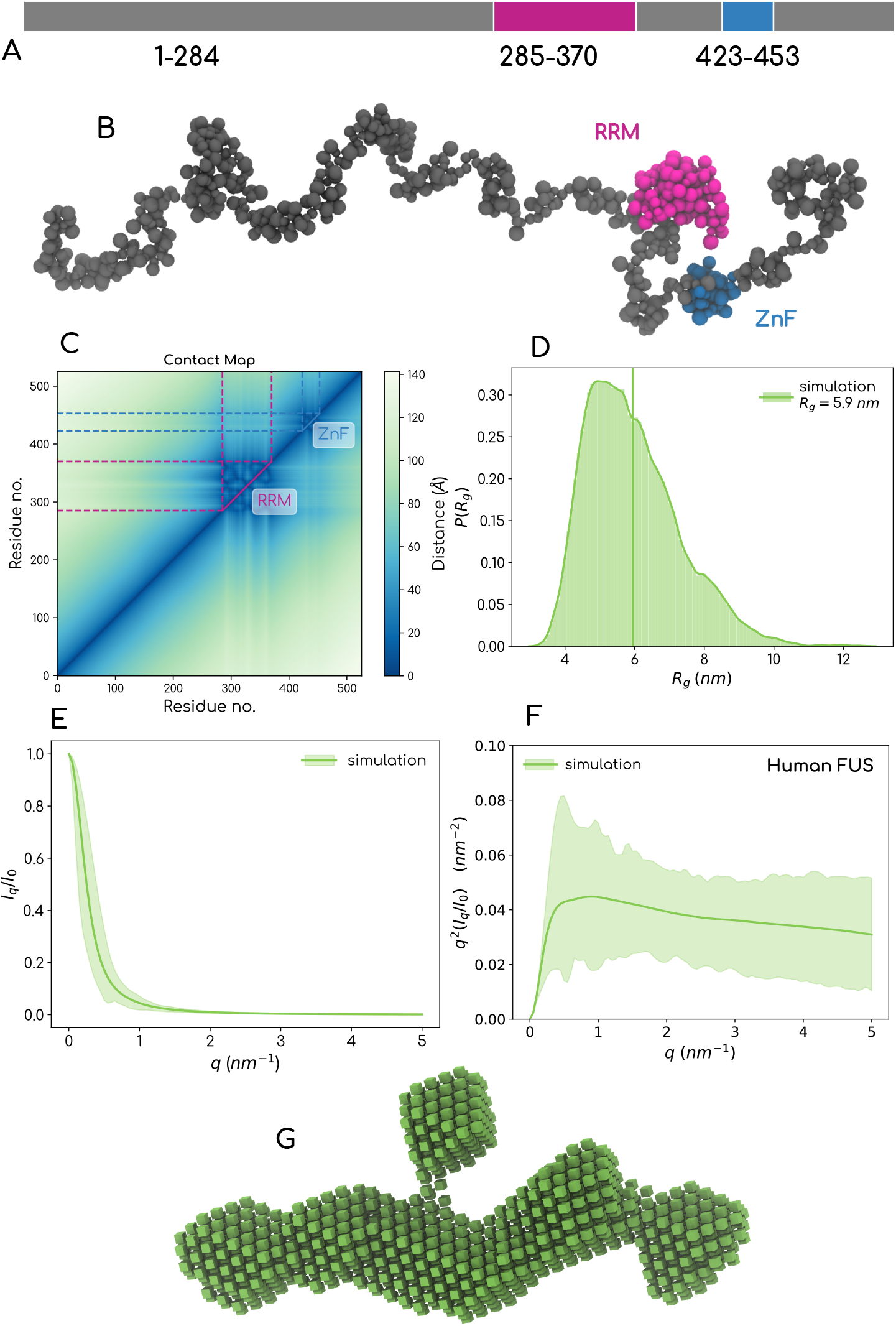
human-FUS : A. Domain architecture B. SOP model C. Contact map D. Radius of Gyration E. Predicted scattering Intensity F. Predicted scattering intensity Kratky representation G. Predicted Scattering envelope of human-FUS

## 4 Conclusions: Scope and Limitations

SOP-MULTI is a MDP CGFF that brings together the SOP-SC and SOP-IDP models with a prescription to model MDF systems with a high level of accuracy with respect to scattering data observed in experiments. We provide this method as an add-on user package that can be compiled with the LAMMPS^54^ engine to generate ensembles of IDPs and MDPs. The model can also be incorporated into other MD engines. Besides the core C++ code in the LAMMPS package, the setup codes written in Python are provided to enable easy setup of the required input files for initiating a LAMMPS simulation of the protein of interest. Through our exhaustive application of this model on multiple proteins, we showed that the model captures experimentally realistic ensembles for 22 IDPs as shown with previous work^28^ as well as works well in generating conformational ensembles of MDPs, keeping the folded regions intact and backbone fluctuations comparable to those in all-atom simulations. We then showed that the single-molecule properties are consistent with data from scattering experiments. A more rigorous approach comparing data from multiple biophysical experiments needs good forward models for individual experimental techniques. We seek to develop such models and comparisons in future studies. One of the observations made from our trajectories is the role of the domain architecture of these proteins in determining the accuracy with which we match the experimental mean. MDPs with long intra-IDRs or IDR tails show relatively more deviation from the experimental mean. This points towards the scope for improving the IDR-folded interactions as a function of salt concentration as these effects get much more pronounced in future applications of this model to study the phase behaviour of these proteins. The model can be used to study much richer biomolecular systems by extending the forcefield definition of SOP with existing CG models of lipid and nucleic acids.

## Supporting information

Supporting Information - User Manual

## Acknowledgement

We thank Prof. Kresten Larsen Lindorff (University of Copenhagen) for the active conversation regarding SAXS and CG models. We thank Dr. Debayan Chakraborty (Institute of Mathematical Sciences, Chennai) for discussions on the self-organized polymer model. We thank Prof. Jeetain Mittal, Texas A&M University, for sharing with us the all-atom trajectory of TDP-43. We thank Kirtika Jha from the Srivastava group for sharing the RMSF data of the HIV-GAG system.

KKB thanks the Ministry of Education, Government of India, for the graduate fellowship and Indo-French Centre for the Promotion of Advanced Research (IFCPAR/CEFIPRA) for the Raman-Charpak fellowship (IFC/4132/RCF 2019/708). AS acknowledges the financial support from the Indian Institute of Science-Bangalore and the high-performance computing facility “Beagle” that was set up from grants by a partnership between the Department of Biotechnology of India and the Indian Institute of Science (IISc-DBT partnership programme). AS thanks the DST for the National Supercomputing Mission grants (DST/NSM/R&D-HPC-Applications/2021/03.10, DST/NSM/RD-HPC-Applications/Extension Grant/2023/27). FIST program sponsored by the Department of Science and Technology and UGC, Centre for Advanced Studies and Ministry of Human Resource Development, India. AS would also like to thank the Teams Science Grant from the DBT-Wellcome Trust India Alliance (Grant number: IA/TSG/21/1/600245). AS also thanks the DBT National Network Project (NNP) grant (BT/PR40323/BTIS/137/78/2023) and the Matrics grants (MTR/2023/001040) from the Science and Engineering Board (SERB), India.

## Author contributions

AS conceived the idea behind the SOP-CGFF for multi-domain proteins. KKB and AS designed the research. KKB and AS wrote the LAMMPS-SOP C++ package. KKB KKB wrote all the pre and post-processing Python tools needed for the package. KKB performed the research and analyzed the data. AS analyzed the data and supervised the study. KKB prepared the first draft of the paper, and KKB and AS polished it together.

## 6 Data availability statement

Our code, models, and curated datasets are publicly available at the laboratory GitHub repository https://github.com/codesrivastavalab/SOP-MULTI.

## References

(1) Jumper, J.; Evans, R.; Pritzel, A.; Green, T.; Figurnov, M.; Ronneberger, O.; Tunyasuvunakool, K.; Bates, R.; Žídek, A.; Potapenko, A., et al. Highly accurate protein structure prediction with AlphaFold. Nature 2021, 596, 583–589.

(2) Subramaniam, S.; Kleywegt, G. J. A paradigm shift in structural biology. Nature Methods 2022, 19, 20–23.

(3) Wei, G.; Xi, W.; Nussinov, R.; Ma, B. Protein ensembles: how does nature harness thermodynamic fluctuations for life? The diverse functional roles of conformational ensembles in the cell. Chemical reviews 2016, 116, 6516–6551.

(4) Thomasen, F. E.; Lindorff-Larsen, K. Conformational ensembles of intrinsically disordered proteins and flexible multidomain proteins. Biochemical Society Transactions 2022, 50, 541–554.

(5) Bhaskara, R. M.; de Brevern, A. G.; Srinivasan, N. Understanding the role of domain– domain linkers in the spatial orientation of domains in multi-domain proteins. Journal of Biomolecular Structure and Dynamics 2013, 31, 1467–1480.

(6) Papaleo, E.; Saladino, G.; Lambrughi, M.; Lindorff-Larsen, K.; Gervasio, F. L.; Nussinov, R. The role of protein loops and linkers in conformational dynamics and allostery. Chemical reviews 2016, 116, 6391–6423.

(7) Ekman, D.; Björklund, Å. K.; Frey-Skött, J.; Elofsson, A. Multi-domain proteins in the three kingdoms of life: orphan domains and other unassigned regions. Journal of molecular biology 2005, 348, 231–243.

(8) Levitt, M. Nature of the protein universe. Proceedings of the National Academy of Sciences 2009, 106, 11079–11084.

(9) Tesei, G.; Trolle, A. I.; Jonsson, N.; Betz, J.; Knudsen, F. E.; Pesce, F.; Johansson, K. E.; Lindorff-Larsen, K. Conformational ensembles of the human intrinsically disordered proteome. Nature 2024, 626, 897–904.

(10) Bashton, M.; Chothia, C. The generation of new protein functions by the combination of domains. Structure 2007, 15, 85–99.

(11) Uversky, V. N.; Oldfield, C. J.; Dunker, A. K. Intrinsically disordered proteins in human diseases: introducing the D2 concept. Annu. Rev. Biophys. 2008, 37, 215–246.

(12) Appadurai, R.; Uversky, V.; Srivastava, A. The Structural and Functional Diversity of Intrinsically Disordered Regions in Transmembrane Proteins. The Journal of Membrane Biology 2019, 252.

(13) Jo, M.; Lee, S.; Jeon, Y.-M.; Kim, S.; Kwon, Y.; Kim, H.-J. The role of TDP-43 propagation in neurodegenerative diseases: integrating insights from clinical and experimental studies. Experimental & molecular medicine 2020, 52, 1652–1662.

(14) Lagier-Tourenne, C.; Polymenidou, M.; Cleveland, D. W. TDP-43 and FUS/TLS: emerging roles in RNA processing and neurodegeneration. Human molecular genetics 2010, 19, R46–R64.

(15) Jean-Philippe, J.; Paz, S.; Caputi, M. hnRNP A1: the Swiss army knife of gene expression. International journal of molecular sciences 2013, 14, 18999–19024.

(16) Wang, B.; Zhang, L.; Dai, T.; Qin, Z.; Lu, H.; Zhang, L.; Zhou, F. Liquid–liquid phase separation in human health and diseases. Signal Transduction and Targeted Therapy 2021, 6, 290.

(17) Wang, J.; Choi, J.-M.; Holehouse, A. S.; Lee, H. O.; Zhang, X.; Jahnel, M.; Maharana, S.; Lemaitre, R.; Pozniakovsky, A.; Drechsel, D., et al. A molecular grammar governing the driving forces for phase separation of prion-like RNA binding proteins. Cell 2018, 174, 688–699.

(18) Martin, E. W.; Thomasen, F. E.; Milkovic, N. M.; Cuneo, M. J.; Grace, C. R.; Nourse, A.; Lindorff-Larsen, K.; Mittag, T. Interplay of folded domains and the disordered low-complexity domain in mediating hnRNPA1 phase separation. Nucleic acids research 2021, 49, 2931–2945.

(19) Kmiecik, S.; Gront, D.; Kolinski, M.; Wieteska, L.; Dawid, A. E.; Kolinski, A. Coarse-grained protein models and their applications. Chemical reviews 2016, 116, 7898–7936.

(20) Kitao, A.; Wagner, G. A space-time structure determination of human CD2 reveals the CD58-binding mode. Proceedings of the National Academy of Sciences 2000, 97, 2064–2068.

(21) Kim, Y. C.; Hummer, G. Coarse-grained models for simulations of multiprotein complexes: application to ubiquitin binding. Journal of molecular biology 2008, 375, 1416–1433.

(22) Kapcha, L. H.; Rossky, P. J. A simple atomic-level hydrophobicity scale reveals protein interfacial structure. Journal of molecular biology 2014, 426, 484–498.

(23) Cragnell, C.; Durand, D.; Cabane, B.; Skepö, M. Coarse-grained modeling of the intrinsically disordered protein Histatin 5 in solution: Monte Carlo simulations in combination with SAXS. Proteins: Structure, Function, and Bioinformatics 2016, 84, 777–791.

(24) Dannenhoffer-Lafage, T.; Best, R. B. A data-driven hydrophobicity scale for predicting liquid–liquid phase separation of proteins. The Journal of Physical Chemistry B 2021, 125, 4046–4056.

(25) Regy, R. M.; Thompson, J.; Kim, Y. C.; Mittal, J. Improved coarse-grained model for studying sequence dependent phase separation of disordered proteins. Protein Science 2021, 30, 1371–1379.

(26) Baul, U.; Chakraborty, D.; Mugnai, M. L.; Straub, J. E.; Thirumalai, D. Sequence effects on size, shape, and structural heterogeneity in intrinsically disordered proteins. The Journal of Physical Chemistry B 2019, 123, 3462–3474.

(27) Baul, U.; Bley, M.; Dzubiella, J. Thermal compaction of disordered and elastin-like polypeptides: a temperature-dependent, sequence-specific coarse-grained simulation model. Biomacromolecules 2020, 21, 3523–3538.

(28) Mugnai, M. L.; Chakraborty, D.; Kumar, A.; Nguyen, H. T.; Zeno, W.; Stachowiak, J. C.; Straub, J. E.; Thirumalai, D. Sizes, conformational fluctuations, and SAXS profiles for Intrinsically Disordered Proteins. bioRxiv 2023, 2023–04.

(29) Joseph, J. A.; Reinhardt, A.; Aguirre, A.; Chew, P. Y.; Russell, K. O.; Espinosa, J. R.; Garaizar, A.; Collepardo-Guevara, R. Physics-driven coarse-grained model for biomolecular phase separation with near-quantitative accuracy. Nature Computational Science 2021, 1, 732–743.

(30) Tesei, G.; Lindorff-Larsen, K. Improved predictions of phase behaviour of intrinsically disordered proteins by tuning the interaction range. Open Research Europe 2023, 2, 94.

(31) Thomasen, F. E.; Pesce, F.; Roesgaard, M. A.; Tesei, G.; Lindorff-Larsen, K. Improving Martini 3 for disordered and multidomain proteins. Journal of Chemical Theory and Computation 2022, 18, 2033–2041.

(32) Fagerberg, E.; Skepo, M. Comparative performance of computer simulation models of intrinsically disordered proteins at different levels of coarse-graining. Journal of Chemical Information and Modeling 2023, 63, 4079–4087.

(33) Seth, S.; Stine, B.; Bhattacharya, A. Fine structures of intrinsically disordered proteins. The Journal of Chemical Physics 2024, 160.

(34) Dignon, G. L.; Zheng, W.; Kim, Y. C.; Best, R. B.; Mittal, J. Sequence determinants of protein phase behavior from a coarse-grained model. PLoS computational biology 2018, 14, e1005941.

(35) Krainer, G.; Welsh, T. J.; Joseph, J. A.; Espinosa, J. R.; Wittmann, S.; de Csilléry, E.; Sridhar, A.; Toprakcioglu, Z.; Gudiškytė, G.; Czekalska, M. A.; Arter, W. E.; Guillén-Boixet, J.; Franzmann, T. M.; Qamar, S.; George-Hyslop, P. S.; Hyman, A. A.; Collepardo-Guevara, R.; Alberti, S.; Knowles, T. P. J. Reentrant liquid condensate phase of proteins is stabilized by hydrophobic and non-ionic interactions. Nature Communications 2021, 12, 1085.

(36) Souza, P. C.; Alessandri, R.; Barnoud, J.; Thallmair, S.; Faustino, I.; Grünewald, F.; Patmanidis, I.; Abdizadeh, H.; Bruininks, B. M.; Wassenaar, T. A., et al. Martini 3: a general purpose force field for coarse-grained molecular dynamics. Nature methods 2021, 18, 382–388.

(37) Periole, X.; Cavalli, M.; Marrink, S.-J.; Ceruso, M. A. Combining an elastic network with a coarse-grained molecular force field: structure, dynamics, and intermolecular recognition. Journal of chemical theory and computation 2009, 5, 2531–2543.

(38) Cao, F.; von Bülow, S.; Tesei, G.; Lindorff-Larsen, K. A coarse-grained model for disordered and multi-domain proteins. bioRxiv 2024, 2024–02.

(39) Thomasen, F. E.; Skaalum, T.; Kumar, A.; Srinivasan, S.; Vanni, S.; Lindorff-Larsen, K. Recalibration of protein interactions in Martini 3. bioRxiv 2023, 2023–05.

(40) Hyeon, C.; Dima, R. I.; Thirumalai, D. Pathways and kinetic barriers in mechanical unfolding and refolding of RNA and proteins. Structure 2006, 14, 1633–1645.

(41) Hyeon, C.; Lorimer, G. H.; Thirumalai, D. Dynamics of allosteric transitions in GroEL. Proceedings of the National Academy of Sciences 2006, 103, 18939–18944.

(42) Liu, Z.; Reddy, G.; Thirumalai, D. Theory of the Molecular Transfer Model for Proteins with Applications to the Folding of the src-SH3 Domain. The Journal of Physical Chemistry B 2012, 116, 6707–6716.

(43) Pincus, D. L.; Thirumalai, D. Crowding effects on the mechanical stability and unfolding pathways of ubiquitin. The journal of physical chemistry B 2009, 113, 359–368.

(44) Liu, Z.; Reddy, G.; O’Brien, E. P.; Thirumalai, D. Collapse kinetics and chevron plots from simulations of denaturant-dependent folding of globular proteins. Proceedings of the National Academy of Sciences 2011, 108, 7787–7792.

(45) Reddy, G.; Liu, Z.; Thirumalai, D. Denaturant-dependent folding of GFP. Proceedings of the National Academy of Sciences 2012, 109, 17832–17838.

(46) Reddy, G.; Thirumalai, D. Collapse precedes folding in denaturant-dependent assembly of ubiquitin. The Journal of Physical Chemistry B 2017, 121, 995–1009.

(47) Mondal, B.; Thirumalai, D.; Reddy, G. Energy landscape of ubiquitin is weakly multidimensional. The Journal of Physical Chemistry B 2021, 125, 8682–8689.

(48) Liu, Z.; Thirumalai, D. Cooperativity and folding kinetics in a multidomain protein with interwoven chain topology. ACS Central Science 2022, 8, 763–774.

(49) Maity, H.; Baidya, L.; Reddy, G. Salt-induced transitions in the conformational ensembles of intrinsically disordered proteins. The Journal of Physical Chemistry B 2022, 126, 5959–5971.

(50) Baidya, L.; Reddy, G. pH Induced Switch in the Conformational Ensemble of Intrinsically Disordered Protein Prothymosin-α and Its Implications for Amyloid Fibril Formation. The Journal of Physical Chemistry Letters 2022, 13, 9589–9598.

(51) Kumar, A.; Chakraborty, D.; Mugnai, M. L.; Straub, J. E.; Thirumalai, D. Sequence determines the switch in the fibril forming regions in the low-complexity FUS protein and its variants. The journal of physical chemistry letters 2021, 12, 9026–9032.

(52) Chakraborty, D.; Straub, J. E.; Thirumalai, D. Differences in the free energies between the excited states of A β 40 and A β 42 monomers encode their aggregation propensities. Proceedings of the National Academy of Sciences 2020, 117, 19926–19937.

(53) Chakraborty, D.; Straub, J. E.; Thirumalai, D. Energy landscapes of Aβ monomers are sculpted in accordance with Ostwald’s rule of stages. Science Advances 2023, 9, eadd6921.

(54) Thompson, A. P.; Aktulga, H. M.; Berger, R.; Bolintineanu, D. S.; Brown, W. M.; Crozier, P. S.; in’t Veld, P. J.; Kohlmeyer, A.; Moore, S. G.; Nguyen, T. D., et al. LAMMPS-a flexible simulation tool for particle-based materials modeling at the atomic, meso, and continuum scales. Computer Physics Communications 2022, 271, 108171.

(55) Van Rossum, G.; Drake Jr, F. L. The python language reference. Python Software Foundation: Wilmington, DE, USA 2014,

(56) Virtanen, P.; Gommers, R.; Oliphant, T. E.; Haberland, M.; Reddy, T.; Cournapeau, D.; Burovski, E.; Peterson, P.; Weckesser, W.; Bright, J., et al. SciPy 1.0: fundamental algorithms for scientific computing in Python. Nature methods 2020, 17, 261–272.

(57) Harris, C. R.; Millman, K. J.; Van Der Walt, S. J.; Gommers, R.; Virtanen, P.; Cournapeau, D.; Wieser, E.; Taylor, J.; Berg, S.; Smith, N. J., et al. Array programming with NumPy. Nature 2020, 585, 357–362.

(58) Kremer, K.; Grest, G. S. Dynamics of entangled linear polymer melts: A moleculardynamics simulation. The Journal of Chemical Physics 1990, 92, 5057–5086.

(59) Betancourt, M. R.; Thirumalai, D. Pair potentials for protein folding: choice of reference states and sensitivity of predicted native states to variations in the interaction schemes. Protein science 1999, 8, 361–369.

(60) Mirdita, M.; Schütze, K.; Moriwaki, Y.; Heo, L.; Ovchinnikov, S.; Steinegger, M. ColabFold: making protein folding accessible to all. Nature methods 2022, 19, 679–682.

(61) Tong, D.; Yang, S.; Lu, L. Accurate optimization of amino acid form factors for computing small-angle X-ray scattering intensity of atomistic protein structures. Journal of Applied Crystallography 2016, 49, 1148–1161.

(62) Rotkiewicz, P.; Skolnick, J. Fast procedure for reconstruction of full-atom protein models from reduced representations. Journal of computational chemistry 2008, 29, 1460–1465.

(63) Svergun, D.; Barberato, C.; Koch, M. H. CRYSOL–a program to evaluate X-ray solution scattering of biological macromolecules from atomic coordinates. Journal of applied crystallography 1995, 28, 768–773.

(64) Stuhrmann, H. B. Interpretation of small-angle scattering functions of dilute solutions and gases. A representation of the structures related to a one-particle scattering function. Acta Crystallographica Section A: Crystal Physics, Diffraction, Theoretical and General Crystallography 1970, 26, 297–306.

(65) Manalastas-Cantos, K.; Konarev, P. V.; Hajizadeh, N. R.; Kikhney, A. G.; Petoukhov, M. V.; Molodenskiy, D. S.; Panjkovich, A.; Mertens, H. D.; Gruzinov, A.; Borges, C., et al. ATSAS 3.0: expanded functionality and new tools for small-angle scattering data analysis. Journal of Applied Crystallography 2021, 54.

(66) Svergun, D. Determination of the regularization parameter in indirect-transform methods using perceptual criteria. Journal of applied crystallography 1992, 25, 495–503.

(67) Hyeon, C.; Dima, R. I.; Thirumalai, D. Size, shape, and flexibility of RNA structures. The Journal of chemical physics 2006, 125.

(68) Fleming, P. J.; Fleming, K. G. HullRad: Fast calculations of folded and disordered protein and nucleic acid hydrodynamic properties. Biophysical journal 2018, 114, 856–869.

(69) Dima, R. I.; Thirumalai, D. Asymmetry in the shapes of folded and denatured states of proteins. The Journal of Physical Chemistry B 2004, 108, 6564–6570.

(70) Vymetal, J.; Vondrasek, J. Gyration-and inertia-tensor-based collective coordinates for metadynamics. Application on the conformational behavior of polyalanine peptides and Trp-cage folding. The Journal of Physical Chemistry A 2011, 115, 11455–11465.

(71) McGibbon, R. T.; Beauchamp, K. A.; Harrigan, M. P.; Klein, C.; Swails, J. M.; Hernández, C. X.; Schwantes, C. R.; Wang, L.-P.; Lane, T. J.; Pande, V. S. MD-Traj: A Modern Open Library for the Analysis of Molecular Dynamics Trajectories. Biophysical Journal 2015, 109, 1528 – 1532.

(72) Eastman, P.; Swails, J.; Chodera, J. D.; McGibbon, R. T.; Zhao, Y.; Beauchamp, K. A.; Wang, L.-P.; Simmonett, A. C.; Harrigan, M. P.; Stern, C. D., et al. OpenMM 7: Rapid development of high performance algorithms for molecular dynamics. PLoS computational biology 2017, 13, e1005659.

(73) Rauscher, S.; Gapsys, V.; Gajda, M. J.; Zweckstetter, M.; De Groot, B. L.; Grubmuller, H. Structural ensembles of intrinsically disordered proteins depend strongly on force field: a comparison to experiment. Journal of chemical theory and computation 2015, 11, 5513–5524.

(74) Fuertes, G.; Banterle, N.; Ruff, K. M.; Chowdhury, A.; Mercadante, D.; Koehler, C.; Kachala, M.; Estrada Girona, G.; Milles, S.; Mishra, A., et al. Decoupling of size and shape fluctuations in heteropolymeric sequences reconciles discrepancies in SAXS vs. FRET measurements. Proceedings of the National Academy of Sciences 2017, 114, E6342–E6351.

(75) Kjaergaard, M.; Nørholm, A.-B.; Hendus-Altenburger, R.; Pedersen, S. F.; Poulsen, F. M.; Kragelund, B. B. Temperature-dependent structural changes in intrinsically disordered proteins: Formation of α–helices or loss of polyproline II? Protein Science 2010, 19, 1555–1564.

(76) Arbesú, M.; Maffei, M.; Cordeiro, T. N.; Teixeira, J. M.; Pérez, Y.; Bernadó, P.; Roche, S.; Pons, M. The unique domain forms a fuzzy intramolecular complex in Src family kinases. Structure 2017, 25, 630–640.

(77) Mittag, T.; Marsh, J.; Grishaev, A.; Orlicky, S.; Lin, H.; Sicheri, F.; Tyers, M.; Forman-Kay, J. D. Structure/function implications in a dynamic complex of the intrinsically disordered Sic1 with the Cdc4 subunit of an SCF ubiquitin ligase. Structure 2010, 18, 494–506.

(78) Wells, M.; Tidow, H.; Rutherford, T. J.; Markwick, P.; Jensen, M. R.; Mylonas, E.; Svergun, D. I.; Blackledge, M.; Fersht, A. R. Structure of tumor suppressor p53 and its intrinsically disordered N-terminal transactivation domain. Proceedings of the National academy of Sciences 2008, 105, 5762–5767.

(79) Johnson, C. L.; Solovyova, A. S.; Hecht, O.; Macdonald, C.; Waller, H.; Grossmann, J. G.; Moore, G. R.; Lakey, J. H. The two-state prehensile tail of the antibacterial toxin colicin N. Biophysical Journal 2017, 113, 1673–1684.

(80) Uversky, V. N.; Gillespie, J. R.; Millett, I. S.; Khodyakova, A. V.; Vasilenko, R. N.; Vasiliev, A. M.; Rodionov, I. L.; Kozlovskaya, G. D.; Dolgikh, D. A.; Fink, A. L., et al. Zn2+-mediated structure formation and compaction of the “natively unfolded” human prothymosin α. Biochemical and Biophysical Research Communications 2000, 267, 663–668.

(81) Lens, Z.; Dewitte, F.; Monté, D.; Baert, J.-L.; Bompard, C.; Sénéchal, M.; Van Lint, C.; De Launoit, Y.; Villeret, V.; Verger, A. Solution structure of the N-terminal transactivation domain of ERM modified by SUMO-1. Biochemical and biophysical research communications 2010, 399, 104–110.

(82) Ahmed, M. C.; Skaanning, L. K.; Jussupow, A.; Newcombe, E. A.; Kragelund, B. B.; Camilloni, C.; Langkilde, A. E.; Lindorff-Larsen, K. Refinement of α-synuclein ensembles against SAXS data: Comparison of force fields and methods. Frontiers in molecular biosciences 2021, 8, 654333.

(83) Holt, C.; Sørensen, E. S.; Clegg, R. A. Role of calcium phosphate nanoclusters in the control of calcification. The FEBS journal 2009, 276, 2308–2323.

(84) Mylonas, E.; Hascher, A.; Bernado, P.; Blackledge, M.; Mandelkow, E.; Svergun, D. I. Domain conformation of tau protein studied by solution small-angle X-ray scattering. Biochemistry 2008, 47, 10345–10353.

(85) Jussupow, A.; Messias, A. C.; Stehle, R.; Geerlof, A.; Solbak, S. M.; Paissoni, C.; Bach, A.; Sattler, M.; Camilloni, C. The dynamics of linear polyubiquitin. Science advances 2020, 6, eabc3786.

(86) Kassem, N.; Araya-Secchi, R.; Bugge, K.; Barclay, A.; Steinocher, H.; Khondker, A.; Wang, Y.; Lenard, A. J.; Bürck, J.; Sahin, C., et al. Order and disorder—An integrative structure of the full-length human growth hormone receptor. Science Advances 2021, 7, eabh3805.

(87) Sonntag, M.; Jagtap, P. K. A.; Simon, B.; Appavou, M.-S.; Geerlof, A.; Stehle, R.; Gabel, F.; Hennig, J.; Sattler, M. Segmental, Domain-Selective Perdeuteration and Small-Angle Neutron Scattering for Structural Analysis of Multi-Domain Proteins. Angewandte Chemie 2017, 129, 9450–9453.

(88) Larsen, A. H.; Wang, Y.; Bottaro, S.; Grudinin, S.; Arleth, L.; Lindorff-Larsen, K. Combining molecular dynamics simulations with small-angle X-ray and neutron scattering data to study multi-domain proteins in solution. PLoS computational biology 2020, 16, e1007870.

(89) Wright, G. S.; Watanabe, T. F.; Amporndanai, K.; Plotkin, S. S.; Cashman, N. R.; Antonyuk, S. V.; Hasnain, S. S. Purification and structural characterization of aggregation-prone human TDP-43 involved in neurodegenerative diseases. Iscience 2020, 23.

(90) Datta, S. A.; Curtis, J. E.; Ratcliff, W.; Clark, P. K.; Crist, R. M.; Lebowitz, J.; Krueger, S.; Rein, A. Conformation of the HIV-1 Gag protein in solution. Journal of molecular biology 2007, 365, 812–824.

(91) Yang, P.; Mathieu, C.; Kolaitis, R.-M.; Zhang, P.; Messing, J.; Yurtsever, U.; Yang, Z.; Wu, J.; Li, Y.; Pan, Q., et al. G3BP1 is a tunable switch that triggers phase separation to assemble stress granules. Cell 2020, 181, 325–345.

(92) Briggs, J. A.; Kräusslich, H.-G. The molecular architecture of HIV. Journal of molecular biology 2011, 410, 491–500.

(93) Ganser-Pornillos, B. K.; Yeager, M.; Sundquist, W. I. The structural biology of HIV assembly. Current opinion in structural biology 2008, 18, 203–217.

(94) Jeffries, C. M.; Ilavsky, J.; Martel, A.; Hinrichs, S.; Meyer, A.; Pedersen, J. S.; Sokolova, A. V.; Svergun, D. I. Small-angle X-ray and neutron scattering. Nature Reviews Methods Primers 2021, 1, 70.

(95) Mohanty, P.; Rizuan, A.; Kim, Y. C.; Fawzi, N. L.; Mittal, J. A complex network of interdomain interactions underlies the conformational ensemble of monomeric TDP-43 and modulates its phase behavior. Protein Science 2024, 33, e4891.

(96) Goldstein, G.; Scheid, M.; Hammerling, U.; Schlesinger, D.; Niall, H.; Boyse, E. Isolation of a polypeptide that has lymphocyte-differentiating properties and is probably represented universally in living cells. Proceedings of the National Academy of Sciences 1975, 72, 11–15.

(97) Komander, D.; Rape, M. The ubiquitin code. Annual review of biochemistry 2012, 81, 203–229.

(98) De Jong, D. H.; Singh, G.; Bennett, W. D.; Arnarez, C.; Wassenaar, T. A.; Schafer, L. V.; Periole, X.; Tieleman, D. P.; Marrink, S. J. Improved parameters for the martini coarse-grained protein force field. Journal of chemical theory and computation 2013, 9, 687–697.

(99) Bonomi, M.; Camilloni, C.; Cavalli, A.; Vendruscolo, M. Metainference: A Bayesian inference method for heterogeneous systems. Science advances 2016, 2, e1501177.

(100) Bonomi, M.; Camilloni, C.; Vendruscolo, M. Metadynamic metainference: Enhanced sampling of the metainference ensemble using metadynamics. Scientific reports 2016, 6, 31232.

(101) Varadi, M.; Anyango, S.; Deshpande, M.; Nair, S.; Natassia, C.; Yordanova, G.; Yuan, D.; Stroe, O.; Wood, G.; Laydon, A., et al. AlphaFold Protein Structure Database: massively expanding the structural coverage of protein-sequence space with high-accuracy models. Nucleic acids research 2022, 50, D439–D444.

(102) Varadi, M.; Bertoni, D.; Magana, P.; Paramval, U.; Pidruchna, I.; Radhakrishnan, M.; Tsenkov, M.; Nair, S.; Mirdita, M.; Yeo, J., et al. AlphaFold Protein Structure Database in 2024: providing structure coverage for over 214 million protein sequences. Nucleic Acids Research 2024, 52, D368–D375.

(103) Mariani, V.; Biasini, M.; Barbato, A.; Schwede, T. lDDT: a local superposition-free score for comparing protein structures and models using distance difference tests. Bioinformatics 2013, 29, 2722–2728.

(104) Martin, E. W.; Peran, I.; Mittag, T. The Collapsed Conformational Landscape of the Hnrnpa1 Low Complexity Region Revealed by SAXS, NMR and Simulation. Biophysical Journal 2018, 114, 367a–368a.

