## Supporting Information - Additional data for "SOP-MULTI: A self-organized polymer based coarse-grained model for multi-domain and intrinsically disordered proteins with conformation ensemble consistent with experimental scattering data"

Table 1: Intrinsically disordered protein systems tested with SOP-MULTI

| | Protein | Length | $R_g^{exp}$ $(nm)$ | $R_g^{sim}$ (nm) | Temperature ($^o$C) | Salt Conc. (mM) |
|---|---|---|---|---|---|---|
| 1 | Histatin5 | 24 | 1.35 | 1.34 | 25 | 150 |
| 2 | RS-repeat | 24 | 1.26 | 1.29 | 25 | 150 |
| 3 | N49 | 36 | 1.69 | 1.57 | 25 | 150 |
| 4 | NLS | 44 | 2.33 | 1.85 | 25 | 150 |
| 5 | ACTR | 71 | 2.63 | 2.34 | 5 | 500 |
| 6 | Nucleoporin | 81 | 2.70 | 2.43 | 25 | 150 |
| 7 | SH4UD | 85 | 2.86 | 2.55 | 25 | 150 |
| 8 | Sic1 | 90 | 3.21 | 2.73 | 25 | 150 |
| 9 | p53 | 93 | 2.87 | 2.77 | 25 | 150 |
| 10 | IBB | 97 | 3.12 | 2.87 | 25 | 150 |
| 11 | CoINT | 98 | 2.83 | 2.72 | 4 | 400 |
| 12 | Prothymosin$-\alpha$ | 111 | 3.78 | 3.78 | 25 | 150 |
| 13 | NUL | 112 | 3.5 | 3.01 | 25 | 150 |
| 14 | ERM | 122 | 3.96 | 3.17 | 25 | 150 |
| 15 | $\alpha-$Synuclein | 140 | 3.50 | 3.45 | 25 | 200 |
| 16 | Osteopontin | 273 | 5.50 | 5.39 | 25 | 80 |
| 17 | K19 | 99 | 3.5 | 2.92 | 15 | 150 |
| 18 | K18 | 130 | 3.8 | 3.46 | 15 | 150 |
| 19 | K17 | 143 | 3.6 | 3.61 | 15 | 150 |
| 20 | K27 | 171 | 3.7 | 3.94 | 15 | 150 |
| 21 | K16 | 174 | 3.9 | 4.1 | 15 | 150 |
| 22 | K32 | 202 | 4.2 | 4.4 | 15 | 150 |

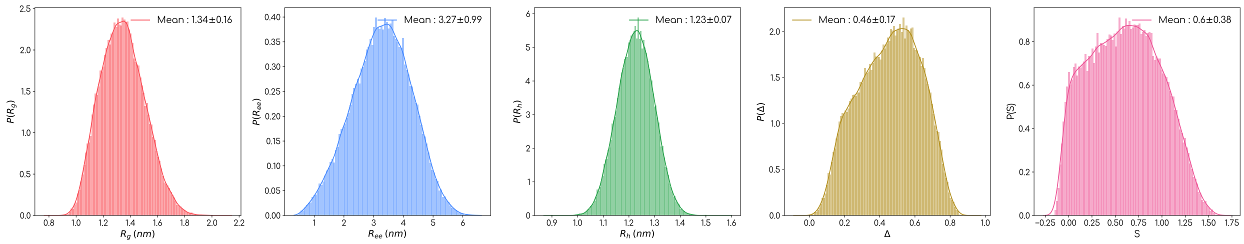


(a) Histatin5

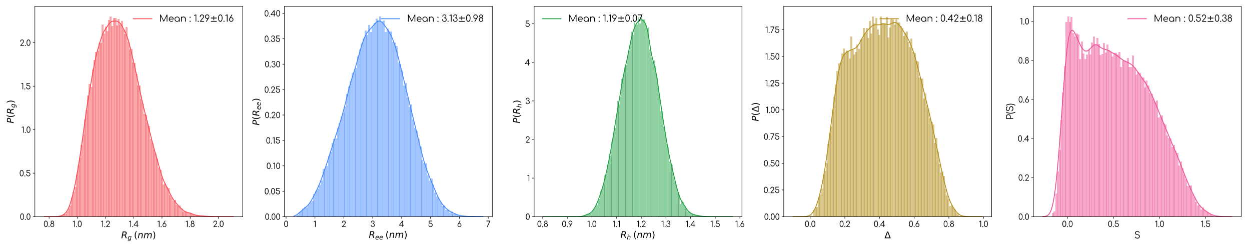


(b) RS-repeat

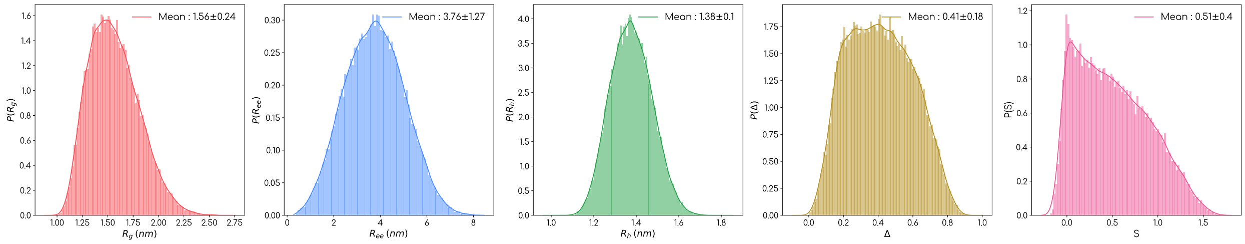


(c) N49

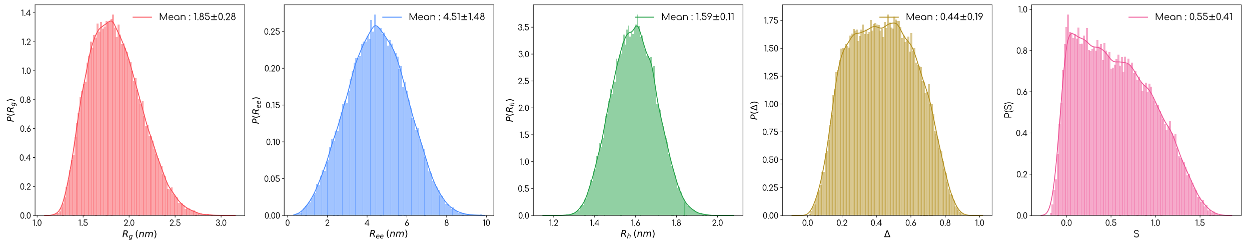


(d) NLS

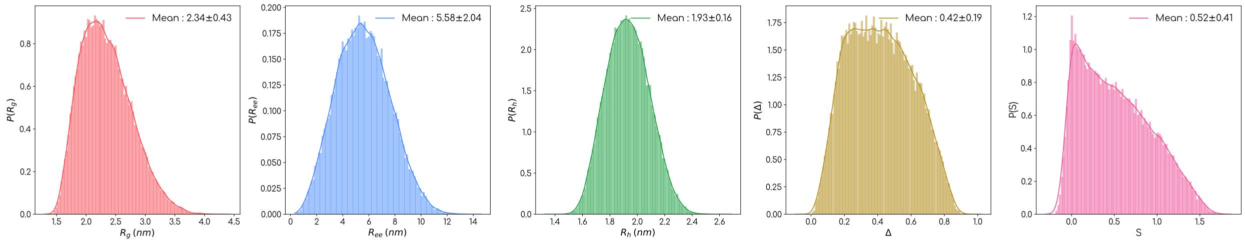


(e) ACTR

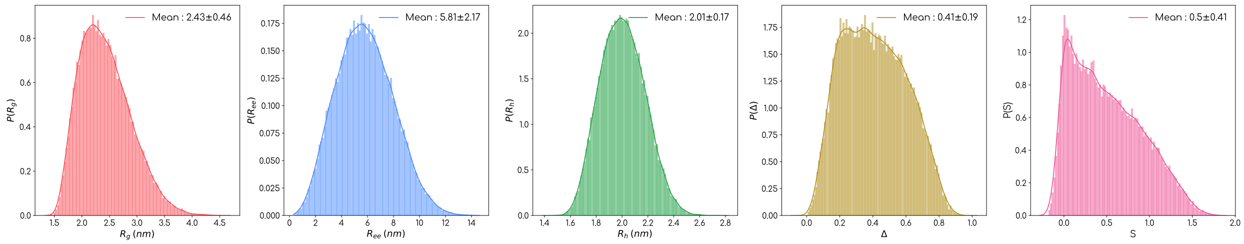


(f) Nucleoporin

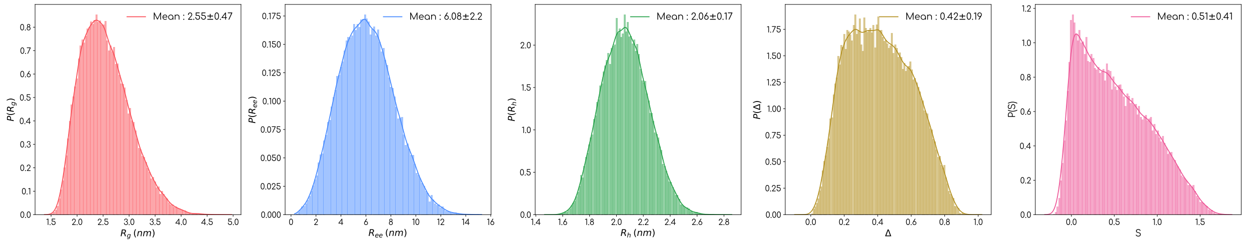


(g) SH4UD

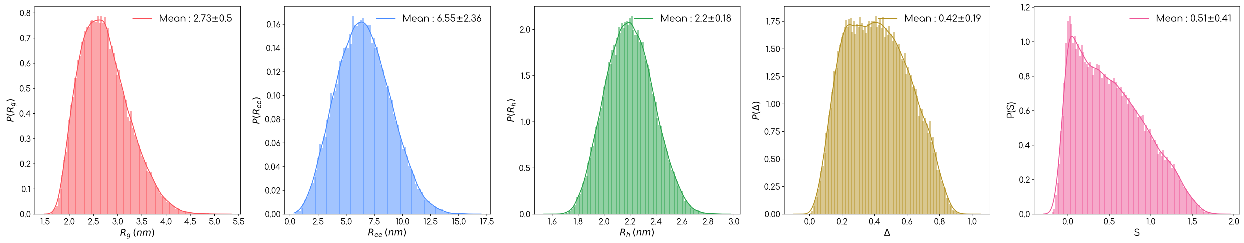


(h) sic1

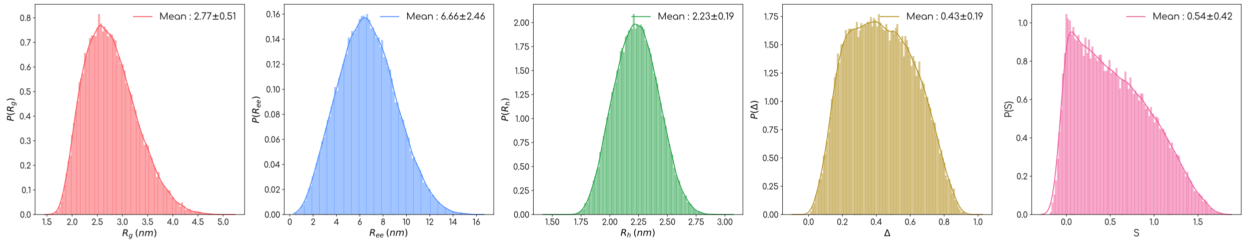


(i) p53

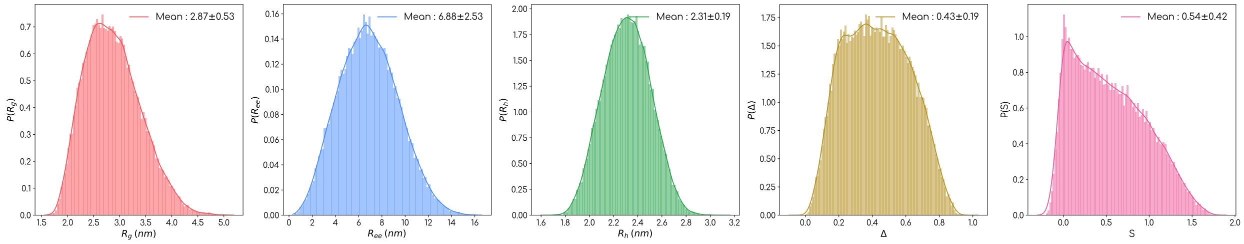


(j) IBB

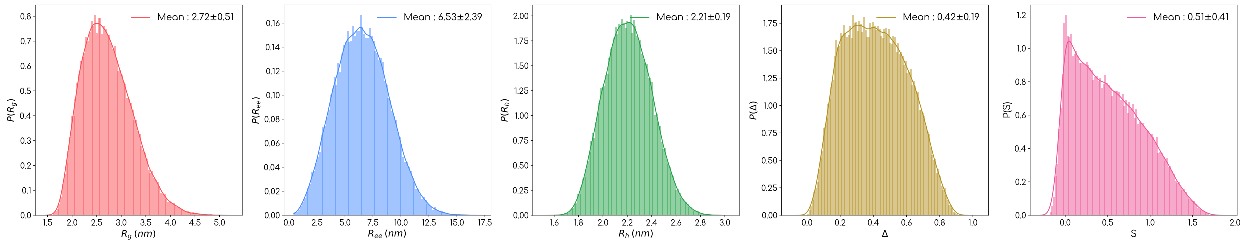


(k) CoINT

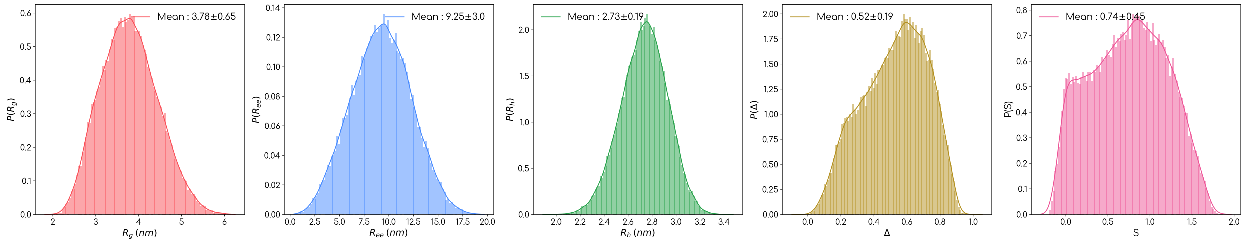


(l) Prothymosin$-\alpha$

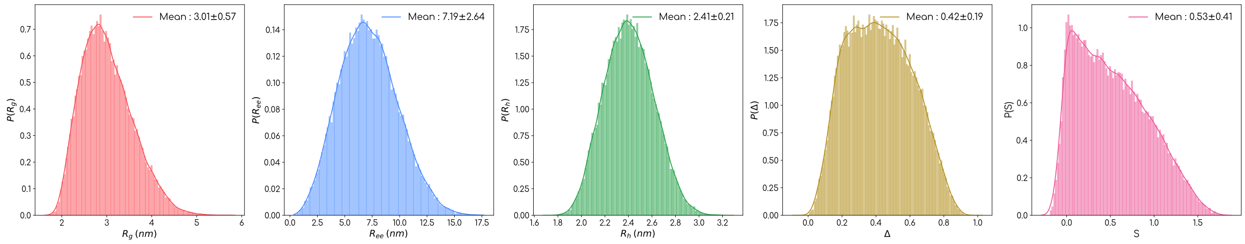


(m) NUL

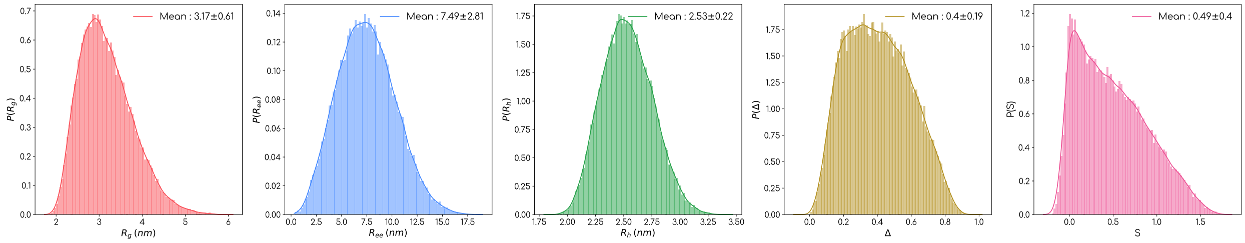


(n) ERM

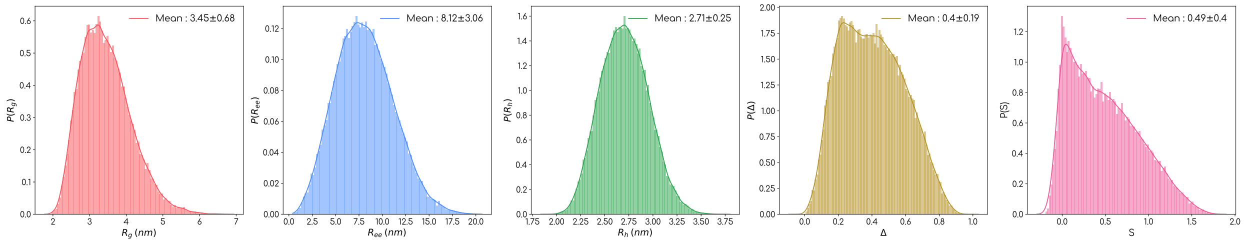


(o) $\alpha-$synuclein

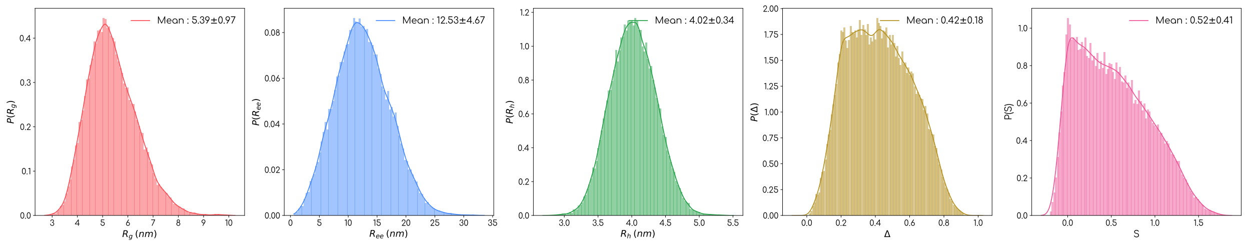


(p) Osteopontin

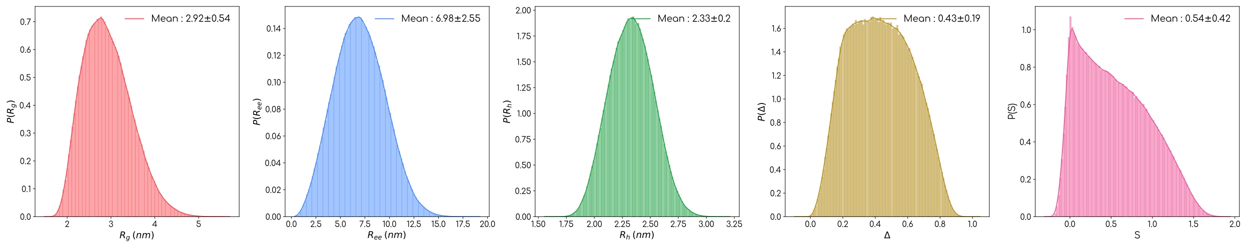


(q) K19

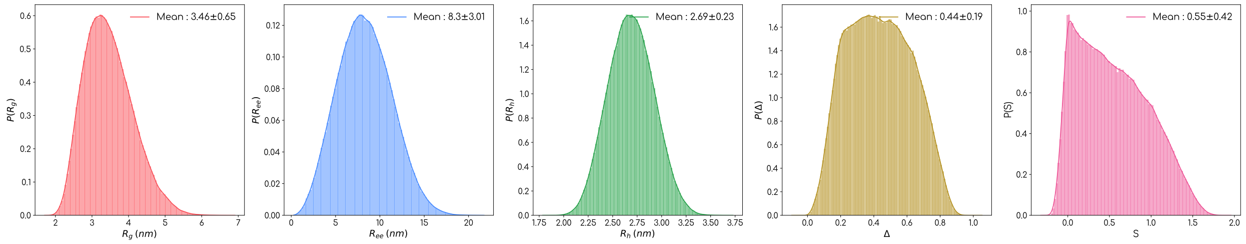


(r) K18

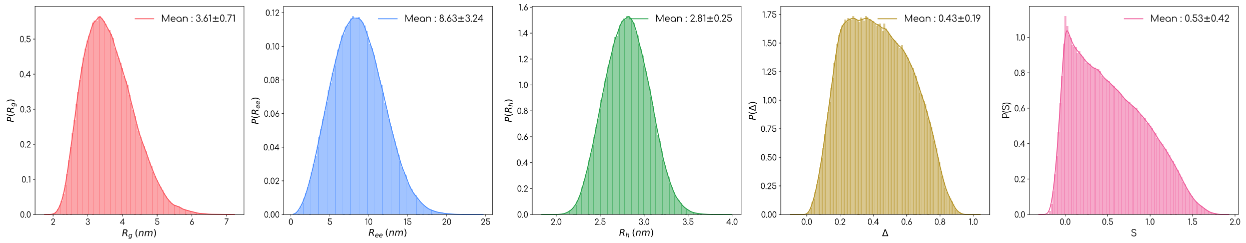


(s) K17

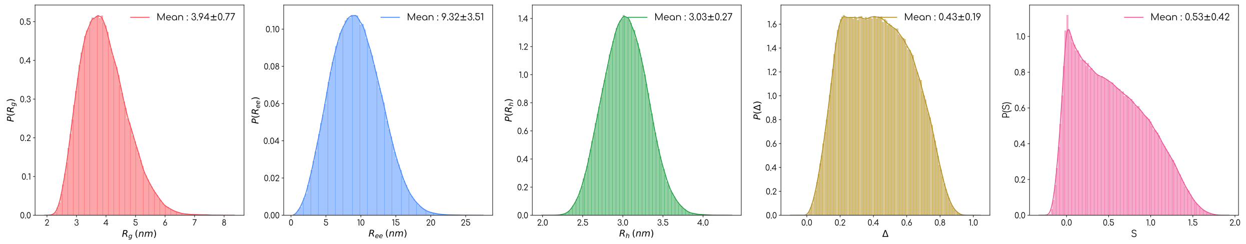


(t) K27

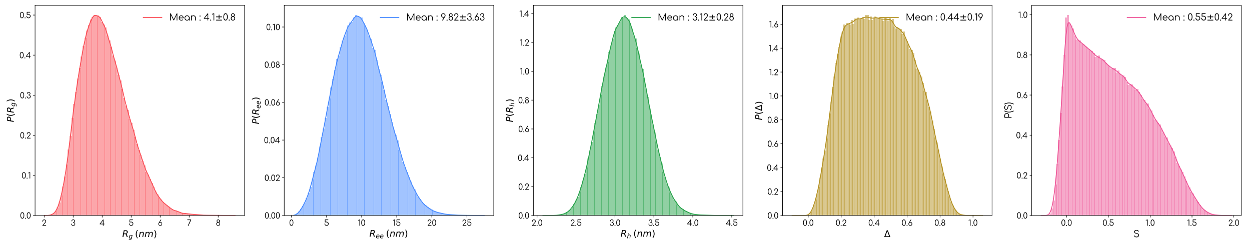


(u) K16

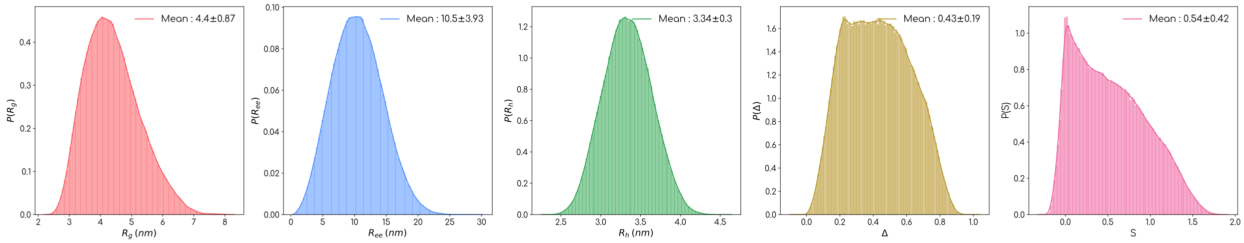


(v) K32

Fig. S 1: Shape properties of IDPs tested with SOP-MULTI package

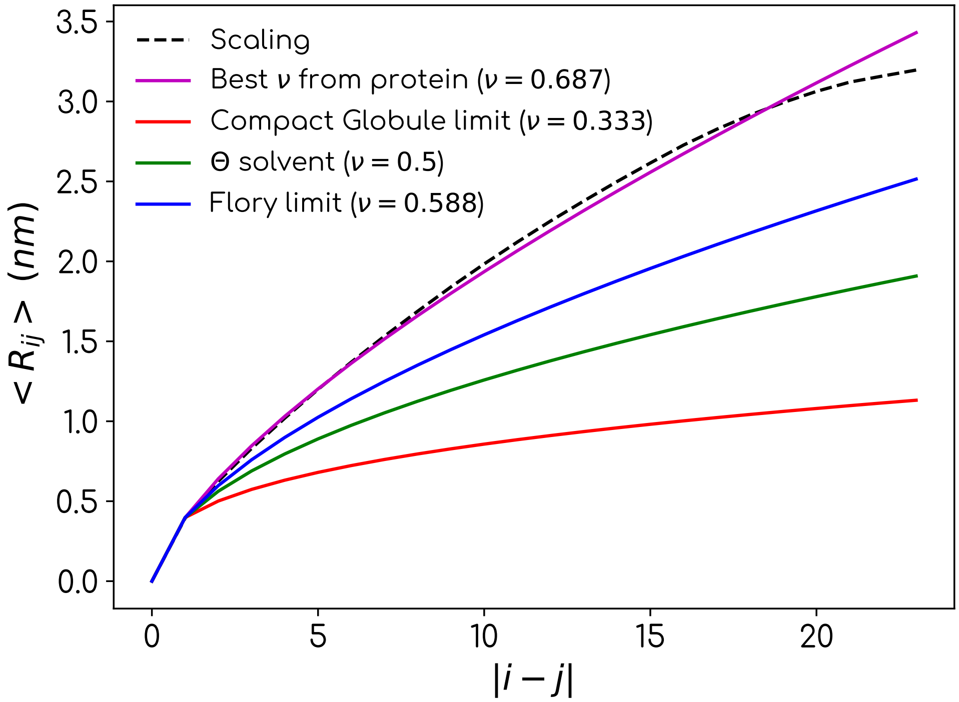


Fig. S 2: Scaling exponent calculation using Histatin5 SOP-MULTI trajectory

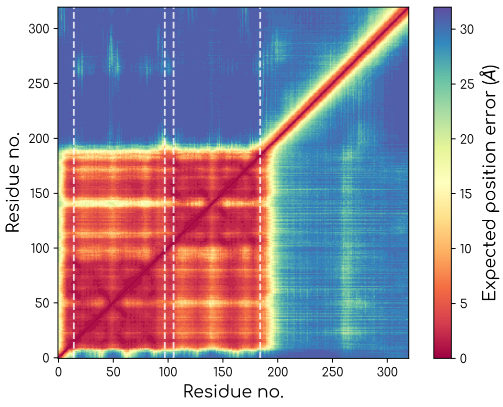


(a) hnrnpa1 PAE

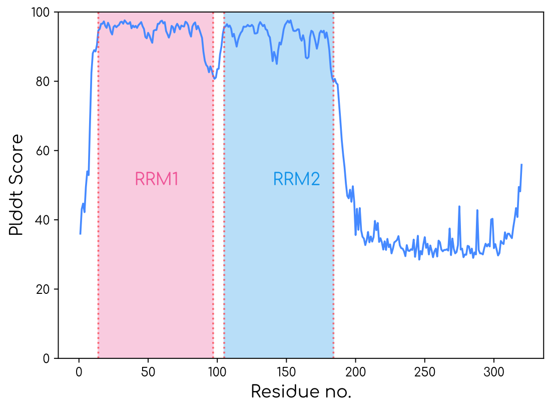


(b) hnrnpa1 PLDDT

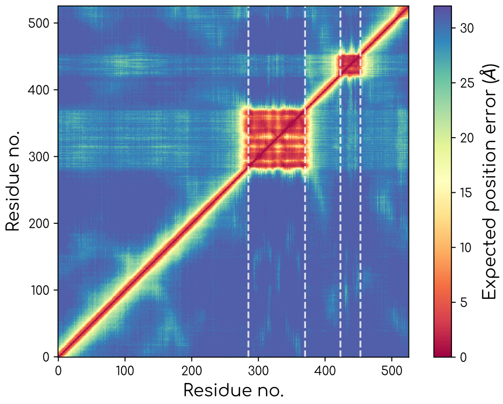


(c) Human FUS PAE

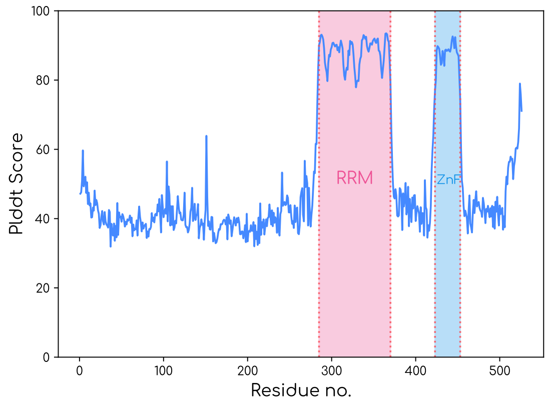


(d) Human FUS PLDDT

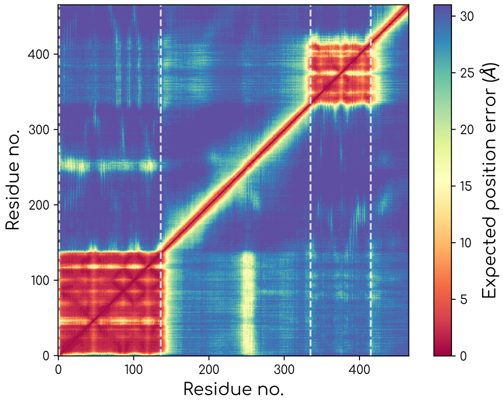


(e) G3BP1 PAE

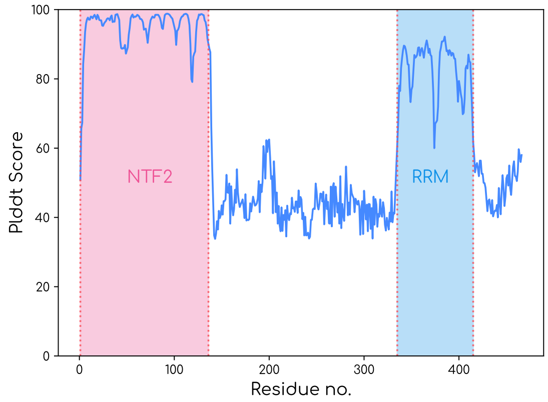


(f) G3BP1 PLDDT

Fig. S3: PAE and PLDDT scores of individual models of multidomain proteins

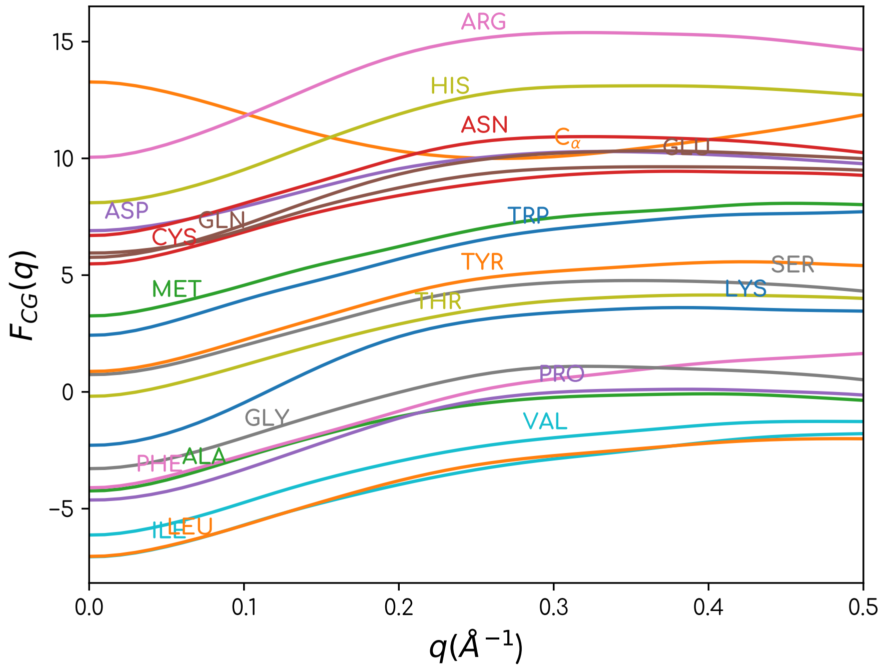


Fig. S 4: Reproduction of CG form factors of two bead model[1]

### References

(1) Tong, D.; Yang, S.; Lu, L. Accurate optimization of amino acid form factors for computing small-angle X-ray scattering intensity of atomistic protein structures. *Journal of Applied Crystallography* **2016**, *49*, 1148–1161.
