## Supporting Information - User Manual for "SOP-MULTI: A self-organized polymer based coarse-grained model for multi-domain and intrinsically disordered proteins with conformation ensemble consistent with experimental scattering data"

This document provides step-by-step instructions for using codes developed as part of the SOP-MULTI package. A description of individual LAMMPS features of SOP-MULTI, along with their syntax and usage examples, are also provided in the last section.

**References 12**

### 1 Compiling LAMMPS executable

A recent version of LAMMPS can be downloaded at the following URL.

https://www.lammps.org/.

Extract the compressed lammps package(.tar.gz) into the destination folder.

Download the SOP-MULTI package of codes from the following Github repository.

https://github.com/krishnakanthbio/SOP-MULTI

Add the files and folders present in "lammps-SOP-MULTI-src" inside the src folder of LAMMPS (replace the files/merge the folders in case of conflict). Open the terminal and change the path to the LAMMPS directory and follow the following instructions to compile the code.

```
$ cd lammps/src   # change to main LAMMPS source folder
$ make yes-SOP-MULTI  # include SOP-MULTI package for compilation
$ make serial     # build a serial LAMMPS executable using GNU g++
$ make mpi        # build a parallel LAMMPS executable with MPI
```

With OPENMP acceleration (Optional*)

```
$ cd lammps/src   # change to main LAMMPS source folder
$ make yes-SOP-MULTI # include SOP-MULTI package for compilation
$ make yes-OPENMP # include OPENMP package for compilation
$ make omp #(needs pre-installed openmpi)
```

*Note : OPENMP acceleration is optional. No significant enhancement in efficiency relative to plain MPI was observed with single-molecule systems.
Based on the installation type, the generated executable file has a filename of **lmp_serial**, **lmp_mpi**, or **lmp_omp**.

### 2 Setting up single-molecule simulation of MDP/IDP in LAMMPS using SOP-MULTI

#### 2.1 Generating all-atom resolution model

An initial all-atom structure modelled using available experimentally solved folded domains and structure prediction tools like Alphafold,[1] iTasser,[2] Robetta,[3] Modeller,[4–6] etc can serve as a good initial configuration for setting up the simulations.

#### 2.2 Generating coarse-grained SOP model

An all-atom structure contains either only heavy atoms or heavy atoms with hydrogens. Based on the type of .pdb file, one can choose the setup code to build a coarse-grained model from the all-atom structure.

**Syntax**

```
$python3 SOP-AA_without_H_2CG-builder <input: all-atom model filename>
<output: coarse-grained model filename>
$python3 SOP-AA_with_H_2CG-builder <input: all-atom model filename>
<output: coarse-grained model filename>
```

**Example**

```
$python3 SOP-AA_without_H_2CG-builder TDP-43-AA.pdb TDP-43-CG.pdb
$python3 SOP-AA_with_H_2CG-builder TDP-43-AA.pdb TDP-43-CG.pdb
```

#### 2.3 Preparing domain information file

The domain information of the protein is provided in the following format in the file named domain_data.dat. This file has three columns. The first and second columns contain the start and end residues of the stretch, and the type of stretch, i.e. IDR or folded, is given in the third column. An IDR is indicated by 0, and folded domains are indicated serially, starting from 1. The absence of this file in the folder models all the beads as type 0, i.e. IDP. An example of such a file for a multidomain protein with three folded domains is shown below.

```
#columns 1:from_residue 2:to_residue 3:domain_id
# 0 indicates IDR/non-folded regions
1 78 1
79 103 0
104 178 2
179 190 0
191 259 3
```

#### 2.4 Generating LAMMPS data file from CG PDB file

The data file to be used with the LAMMPS executable can be generated using the CG .pdb file and domain_data.dat file(not needed for IDPs) generated from the previous steps.

**Syntax**

```
$python3 SOP-PDB2data.py <input: coarse-grained model filename>
<output: LAMMPS data filename>
```

**Example**

```
$python3 SOP-PDB2data.py TDP-43-CG.pdb data.TDP-43
```

#### 2.5 Generating lists of bead pairs in folded domains

The data file to be used with the LAMMPS executable can be generated using the CG .pdb file generated in the previous step.

**Syntax**

```
$python3 SOP-data2pair_list.py <input: LAMMPS data filename>
<output: pairs list filename>
```

**Example**

```
$python3 SOP-data2pair_list.py data.TDP-43 folded_pair_list.txt
```

#### 2.6 Generating forcefield file

The forcefield file to be used for simulating MDP/IDP can be generated by using the following code. The inputs are Temperature($^oC$) and Salt Concentration($mM$)

**Syntax**

```
$python3 ff_writer.py <input: LAMMPS data filename>
<output: pairs list filename>
```

**Example**

```
$python3 ff_writer.py 25 150
```

#### 2.7 Preparing LAMMPS settings file and running the simulation

The LAMMPS input file is the primary file that is provided as input to the LAMMPS executable

```
$(path to executable)/lmp_serial -i in.sop_multi # on single CPU
$mpirun (path to executable)/lmp_mpi -i in.sop_multi # on multiple CPUs
```

An example script for preparing a SOP-MULTI compatible LAMMPS input file can be found at the following weblink
https://github.com/krishnakanthbio/SOP-MULTI

### 3 Components of SOP-MULTI package

#### 3.1 *sop_bead* atom style

The commonly used atom style in LAMMPS[7] for bio-molecules is the *full* atom style[8] which accommodates bonds, angles, dihedrals, impropers and charge. However, in the case of the SOP-MULTI model, the LAMMPS code should recognize which domain and residue of the protein each bead belongs to. We added a new atom style named *sop_bead*, which allows bead's domain and residue information in the input file. A comparison of *full* with *sop_bead* atom style is shown below.

```
full atom style

atom-id molecule-tag atom-type q x y z

sop_bead atom style

atom-id molecule-tag domain-tag residue-tag atom-type q x y z
```

**Usage:**

```
atom_style sop_bead
```

#### 3.2 *sop_ff* special bonds

The self-organized polymer model uses an excluded volume potential to deal with all 1-2 and 1-3 interactions. The default neighbour list of LAMMPS includes interactions 1-4 and above. These pairs are made accessible to the neighbour list by setting the corresponding coefficients to one using **special_bonds**.

**Usage:**

```
special_bonds sop_ff
```

#### 3.3 bond_style *fene/sop*

The bonded interaction is modelled using a Fene Potential. The pre-existing code available for fene[9,10] bond in the MOLECULE package of LAMMPS was modified for use with SOP-MULTI model.

$$E = -\frac{K}{2} R_0^2 log\left(1 - \frac{(r - r_{ref})^2}{R_0^2}\right)$$

**Syntax**

```
bond_style fene/sop
```

**Example**

```
bond_style fene/sop
bond_coeff 1 20.0 2.0 3.6018
```

The following coefficients are assigned to each bond type using the bond_coeff command as in the example above, or in the "Bond Coeffs" section of data file:

- $K$ (energy/distance$^2$)
- $R_0$ (distance)
- $r_{ref}$ (distance)

#### 3.4 pair_style *debye/sop*

The *debye/sop* style computes the Coulombic pairwise interaction between charged beads within the provided cutoff $r_c$ with an exponential decaying damping factor to mimic the screening effect of a polar solvent.[11] $\kappa$ is the inverse of the Debye length and $r_c$ is the cutoff. This potential is inactive between all the charged beads belonging to the same domain and beads separated by 1-2 and 1-3 connectivity. The dielectric constant can be modified using the LAMMPS keyword **dielectric**.

$$E = \epsilon_{debye} \frac{q_i q_j exp(-\kappa r)}{\varepsilon r} \quad r < r_c$$

**Syntax**

```
dielectric value
pair_style debye/sop global_cutoff
pair_coeff debye/sop epsilon_debye kappa local_cutoff
```

**Example**

```
dielectric 77.505
pair_style debye/sop 35.0
pair_coeff 4 5 debye/sop 1.0 0.1806 35.0
```

- epsilon_debye = prefactor of electrostatic interaction (energy units)
- kappa = Debye length (inverse distance units)
- cutoff = global cutoff (distance units)

#### 3.5 pair_style *lj/sop*

The pair_style *lj/sop* encodes Lennard-Jones potential used in SOP-IDP[12] model along with excluded volume between non-bonded beads.

If the pair of beads are connected by 1-2 or 1-3 connectivity, then the excluded volume potential($E_{exc}$) is activated. All the pairs with 1-4 and above connectivity and within cutoff $r_c$ are dealt by LJ potential($E_{LJ}$). However this LJ potential is inactive inside individual folded domains.

$$E_{LJ} = \epsilon_{LJ}\left[\left(\frac{\sigma}{r}\right)^{12} - 2\left(\frac{\sigma}{r}\right)^{6}\right] \quad r < r_c$$

$$E_{exc} = \epsilon_{exc}\left(\frac{\sigma}{r}\right)^{6}$$

**Syntax**

```
pair_style lj/sop cutoff
pair_style lj/sop epsilon_excluded epsilon_LJ sigma
```

**Examples**

```
pair_style lj/sop 25.0
pair_coeff 1 2 lj/sop 1.0 0.13697 4.15
```

- $\epsilon_{exc}$ (energy units)
- $\epsilon_{lj}$ (energy units)
- $\sigma$ (distance units)

#### 3.6 pair_style *list/sop*

The folded domains of the protein molecules use a pre-defined pair list of beads. The existing pair_style list command[13] was extended to accommodate the Lennard-Jones potential of the Self-Organized Polymer(SOP) Model. The pair style *list/sop* computes interactions between explicitly listed pairs of beads. Because the parameters are set in the list file, the pair_coeff command has no parameters (but still needs to be provided). The check and nocheck keywords enable/disable tests that check whether all listed pairs of atom IDs were present and the interactions computed. If nocheck is set and either bead ID is not present, the interaction is skipped.

**Syntax**

```
pair_style list/sop listfile cutoff keyword
```

- listfile = name of the file with list of pairwise interactions
- cutoff = global cutoff (distance units)
- keyword = optional flag *nocheck* or *check* (default is *check*)

**Example**

```
pair_style list/sop folded_pair_list.txt 200.0
pair_coeff * *

pair_style hybrid/overlay lj/sop 25.0 debye/sop 35.0 list/sop folded_pair_list.txt
20.0
pair_coeff * * list/sop
```

The format of the list file is as follows:

- one line per pair of atoms
- empty lines will be ignored
- comment text starts with a '#' character
- line syntax: *ID1 ID2 style coeffs cutoff*

```
ID1 = atom ID of first atom
ID2 = atom ID of second atom
style = style of interaction
coeffs = list of coeffs
cutoff = cutoff for interaction (optional)
```

Here is an example file:

```
# folded_pair_list.txt
487 493 ljsop 0.45 3.8
487 494 ljsop 0.18 4.83
487 495 ljsop 0.45 3.8
487 496 ljsop 0.18 4.91
487 497 ljsop 0.45 4.15
```

The style *lj/sop* computes pairwise interactions with the formula

$$E = \epsilon\left[\left(\frac{r_i^{fold}}{r}\right)^{12} - 2\left(\frac{r_i^{fold}}{r}\right)^{6}\right]$$

- $\epsilon$ (energy units)
- $r^{fold}$ (distance units)

### References

(1) Jumper, J.; Evans, R.; Pritzel, A.; Green, T.; Figurnov, M.; Ronneberger, O.; Tunyasuvunakool, K.; Bates, R.; Žídek, A.; Potapenko, A., et al. Highly accurate protein structure prediction with AlphaFold. *Nature* **2021**, *596*, 583–589.

(2) Zhou, X.; Zheng, W.; Li, Y.; Pearce, R.; Zhang, C.; Bell, E. W.; Zhang, G.; Zhang, Y. I-TASSER-MTD: a deep-learning-based platform for multi-domain protein structure and function prediction. *Nature Protocols* **2022**, *17*, 2326–2353.

(3) Baek, M.; DiMaio, F.; Anishchenko, I.; Dauparas, J.; Ovchinnikov, S.; Lee, G. R.; Wang, J.; Cong, Q.; Kinch, L. N.; Schaeffer, R. D., et al. Accurate prediction of protein

structures and interactions using a three-track neural network. *Science* **2021**, *373*, 871–876.

(4) Šali, A.; Blundell, T. L. Comparative protein modelling by satisfaction of spatial restraints. *Journal of molecular biology* **1993**, *234*, 779–815.

(5) Fiser, A.; Do, R. K. G.; Šali, A. Modeling of loops in protein structures. *Protein science* **2000**, *9*, 1753–1773.

(6) Webb, B.; Sali, A. Comparative protein structure modeling using MODELLER. *Current protocols in bioinformatics* **2016**, *54*, 5–6.

(7) Thompson, A. P.; Aktulga, H. M.; Berger, R.; Bolintineanu, D. S.; Brown, W. M.; Crozier, P. S.; in't Veld, P. J.; Kohlmeyer, A.; Moore, S. G.; Nguyen, T. D., et al. LAMMPS-a flexible simulation tool for particle-based materials modeling at the atomic, meso, and continuum scales. *Computer Physics Communications* **2022**, *271*, 108171.

(8) atom_style command. 2023; `https://docs.lammps.org/atom_style.html`.

(9) Kremer, K.; Grest, G. S. Dynamics of entangled linear polymer melts: A molecular-dynamics simulation. *The Journal of Chemical Physics* **1990**, *92*, 5057–5086.

(10) bond_style fene/expand command. 2023; `https://docs.lammps.org/bond_fene_expand.html`.

(11) pair_style pair_style lj/cut/coul/debye command. 2023; `https://docs.lammps.org/pair_lj_cut_coul.html#pair-style-lj-cut-coul-debye-command`.

(12) Baul, U.; Chakraborty, D.; Mugnai, M. L.; Straub, J. E.; Thirumalai, D. Sequence effects on size, shape, and structural heterogeneity in intrinsically disordered proteins. *The Journal of Physical Chemistry B* **2019**, *123*, 3462–3474.

(13) pair_style list command. 2023; `https://docs.lammps.org/pair_list.html`.
